# DNA remodeling couples target recognition to directional transposition in a Tn7-like CAST

**DOI:** 10.64898/2026.06.02.729338

**Authors:** Shukun Wang, Romana Siddique, Leifu Chang

## Abstract

CRISPR-associated transposons (CASTs) couple target recognition to the insertion of large DNA cargoes, but how distinct targeting pathways are converted into productive and directional integration remains poorly understood^1^. Here we define the assembly pathway of a type I-B1 CAST from *Anabaena variabilis*, a system closely related to prototypical Tn7 that retains TnsD-mediated *glmS* recognition while incorporating CRISPR-based RNA-guided targeting^2^. Cryo-electron microscopy structures across multiple intermediates define a structural trajectory from two-step TnsD-mediated target recognition and stepwise assembly of the AAA+ ATPase TnsC to recruitment and activation of the split TnsA/TnsB transposase module. This trajectory culminates in an asymmetric strand-transfer complex that provides a structural basis for insertion orientation and supports a role for ATP hydrolysis in productive transpososome assembly. In parallel, RNA-guided targeting structures refine the functional PAM to ATG, define TniQ recruitment by Cascade, and show how CRISPR-based recognition converges on the shared TnsABC machinery. Together, these findings establish DNA remodeling and minor-groove positioning of TnsC as a common structural signal that converts protein- and RNA-guided target recognition into directional DNA insertion.

## Introduction

Precise insertion of large DNA segments by DNA transposons requires target recognition to be tightly coordinated with transpososome assembly and strand transfer. The bacterial transposon Tn7 provides a classic example of site-specific DNA integration, using the protein-guided target selector TnsD to recognize the chromosomal site at the 3′ end of *glmS*^3,4^. A subset of Tn7-like transposons has co-opted CRISPR–Cas systems^5–7^, giving rise to CRISPR-associated transposons (CASTs) that couple RNA-guided target recognition to DNA insertion^2,8–12^. These systems offer a natural solution to a problem that has been difficult to solve by engineering alone: how to connect programmable target selection to productive, directional integration of large DNA cargoes. Recent advances have extended CAST-mediated DNA insertion to mammalian cells, highlighting their promise as programmable platforms for large-cargo genome engineering^13–15^. CAST systems are mechanistically diverse^1,16^. Type V-K systems use Cas12k, TniQ and the host factor S15 to recruit filamentous TnsC assemblies and activate a TnsB transposase^17–22^, whereas type I CASTs use Cascade-like effectors and TniQ to engage ring-like TnsC assemblies that activate the TnsA/TnsB transposase^23–30^. Even within type I CASTs, independent evolutionary acquisitions of CRISPR–Cas modules have produced subtypes with distinct architectures and coupling mechanisms. Type I-B2 systems encode fused TnsAB proteins^2,23,24^, whereas type I-B1 and type I-F systems encode separate TnsA and TnsB proteins^2,8^. Some Tn7-like CASTs (e.g., I-B1, I-B2^2^, and I-D^10^) retain TnsD-mediated chromosomal targeting while also using CRISPR effector complexes and TniQ for RNA-guided targeting. Furthermore, structural studies of prototypical Tn7 have revealed mechanistic diversity in TnsD-dependent targeting^31–35^, including stepwise TnsC oligomerization^34^ and TnsA recruitment^32^, highlighting a regulatory logic distinct from previously characterized CAST systems. This diversity has made it difficult to infer, from any single subtype, how target recognition is converted into productive DNA insertion.

A key unresolved question is whether target DNA serves merely as a binding platform or, through structural remodeling, also provides a signal that promotes transpososome assembly. In this study, we address this question using the type I-B1 CAST from *Anabaena variabilis* (*Av*CAST), a system closely related to prototypical Tn7 that retains a canonical TnsD-guided *glmS* targeting pathway, as in Tn7^36,37^, as well as an RNA-guided pathway^2,10^. Using biochemical reconstitution and cryo-electron microscopy, we determined structures representing multiple stages of both *Av*CAST targeting pathways. These structures define a trajectory from two-step TnsD-mediated target recognition and stepwise TnsC assembly to recruitment and activation of the split TnsA/TnsB transposase module. The pathway leads to an asymmetric strand-transfer complex (STC) that provides a structural basis for insertion orientation and supports a role for ATP hydrolysis in productive transpososome assembly. In parallel, structures of *Av*Cascade–DNA, *Av*Cascade–TniQ–DNA and TniQ–TnsC–DNA assemblies explain how RNA-guided targeting is connected to the same downstream machinery. In both pathways, target recognition is coupled to minor-groove positioning of TnsC on target DNA, suggesting that DNA geometry acts as a shared structural signal that licenses productive transpososome assembly. Together, our data define how a Tn7-like CAST integrates protein- and RNA-guided targeting into a common assembly program for directional DNA insertion.

## Results

### Biochemical reconstitution of dual targeting pathways in *Av*CAST

The *Av*CAST locus encodes a type I-B CRISPR effector complex, a CRISPR array, core transposition proteins and associated cargo genes^2^ **(Fig. 1a)**. To enable structural and mechanistic dissection, we biochemically reconstituted *Av*CAST using purified TnsA, TnsB, TnsC, TnsD, TniQ and *Av*Cascade, together with donor and target plasmids designed to monitor both TnsD-guided and RNA-guided transposition **(Extended Data Fig. 1a,b and Supplementary Fig. 1)**. The target plasmid contains two recognition sites: a TnsD target site comprising the last 50 bp of the *glmS* coding sequence, including the stop codon, and an *Av*Cascade target site comprising a protospacer preceded by a 5′-ATG-3′ PAM, as defined below. The donor plasmid contains the left and right transposon ends (LE and RE) flanking a kanamycin-resistance cassette **(Extended Data Fig. 1b)**.

**Fig. 1.**
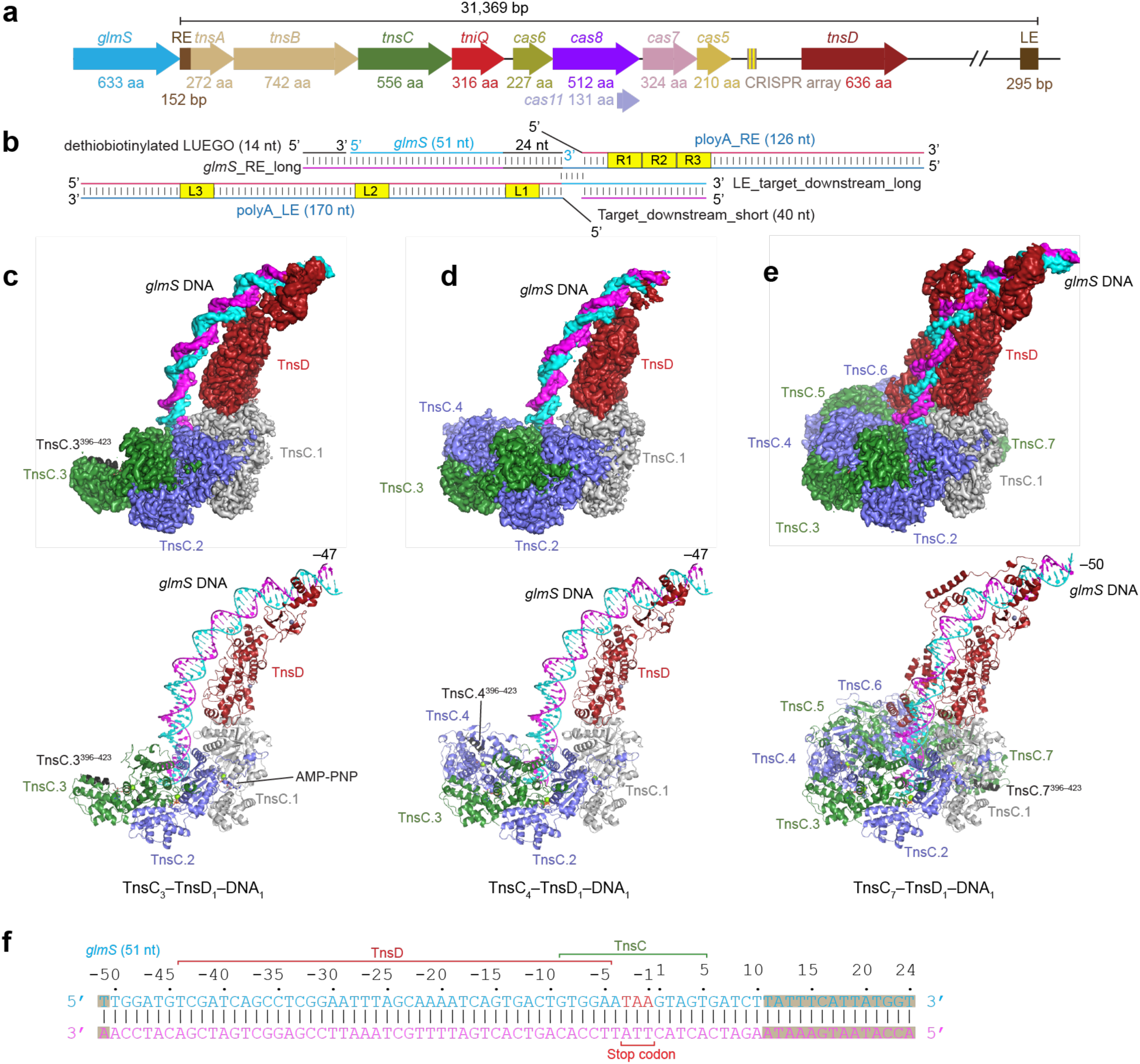
Cryo-EM structures of TnsD–TnsC assemblies on *glmS* DNA. **a**, Genetic organization of the *Av*CAST locus, showing the *glmS* target gene, transposon ends, transposition genes, CRISPR–Cas genes and CRISPR array. **b**, Schematic of the desthiobiotinylated DNA substrate used for assembly, containing the *glmS* target sequence, downstream target DNA, and joined LE and RE transposon ends. TnsB-binding sites on LE and RE are indicated. **c–e**, Cryo-EM density maps and atomic models of the TnsC_3_–TnsD_1_–DNA_1_, TnsC_4_–TnsD_1_–DNA_1_ and TnsC_7_–TnsD_1_–DNA_1_ assemblies, respectively. **f**, Schematic of the modeled *glmS* target DNA in TnsC_7_-TnsD_1_-DNA_1_, showing regions contacted by TnsD and TnsC. The *glmS* stop codon is indicated. Shaded regions indicate nucleotides that were not modeled in the structure.

After incubation of purified proteins with donor and target plasmids in the presence of ATP and Mg^2+^, transposition products were detected by direct PCR, qPCR and BamHI cleavage assays **(Extended Data Fig. 1c–n)**. In the TnsD-guided pathway, sequencing of recovered plasmids showed that 11 of 12 products contained precise insertions 24 bp downstream of the *glmS* stop codon, with RE consistently positioned proximal to the target site **(Extended Data Fig. 1f,j)**. RE contains three contiguous TnsB-binding sites, whereas LE contains three more widely spaced TnsB-binding sites. Truncation assays using shortened LE and RE substrates further defined the minimal requirements for transposition: the end-proximal TnsB-binding site on LE and all three TnsB-binding sites on RE were required for detectable activity **(Extended Data Fig. 1h,i and Extended Data Fig. 2a,b)**. Replacing ATP with the non-hydrolyzable analogue AMP-PNP markedly reduced transposition efficiency, from approximately 7.6% to 0.6%, but still yielded detectable on-target insertions compared with the no-ATP control **(Extended Data Fig. 1d)**. Among products generated in the presence of AMP-PNP, 10 of 12 retained insertion at the canonical position 24 bp downstream of *glmS* **(Extended Data Fig. 1g)**. These results indicate that ATP hydrolysis promotes productive transposition without substantially altering TnsD-guided insertion-site selection. In the RNA-guided pathway, sequencing of recovered plasmids showed that all 12 products were on target, with insertions distributed within a narrow window 78–81 bp downstream of the PAM **(Extended Data Fig. 1k–n)**, consistent with previous studies of *Av*CAST in *E. coli* ^2^.

### Cryo-EM structures of TnsD–TnsC assemblies on *glmS* DNA

We first sought to assemble a TnsABCD transpososome complex using a desthiobiotinylated DNA substrate in the presence of AMP-PNP. Guided by structures of STCs from Mu^38^, the P element^39^, type V-K CAST^20–22^ and type I-B2 CAST^24^, we designed the substrate to mimic a post-transposition product in which the LE and RE were joined 24 bp downstream of the *glmS* target sequence. The target region contained 51 bp of the *glmS* coding sequence followed by 40 bp of downstream DNA (**Fig. 1b**)

Although SDS-PAGE analysis of the eluted fraction revealed clear bands corresponding to TnsA, TnsB, TnsC, and TnsD (**Extended Data Fig. 2c**), cryo-EM analysis identified particles corresponding to TnsC–TnsD–DNA complexes exclusively. Further analysis revealed three distinct stoichiometric assemblies: TnsC_3_–TnsD_1_–DNA_1_ (55.3%), TnsC_4_–TnsD_1_–DNA_1_ (6.8%) and TnsC_7_–TnsD_1_–DNA_1_ (37.9%) (**Fig. 1c–e, Extended Data Fig. 2d,e and 3, Supplementary Fig. 2 and Extended Data Table 1**), determined at resolutions of 3.08 Å, 3.82 Å, and 3.17 Å, respectively. In contrast, when the same assembly was performed with ATP instead of AMP-PNP, TnsC–TnsD–DNA complexes were no longer detected, and a STC containing TnsA, TnsB and the C-terminal region of TnsC was observed **(Extended Data Fig. 2f–h and 4, and Extended Data Table 1)**. These results indicate that the non-hydrolyzable ATP analogue stabilizes target-recognition assemblies, whereas ATP supports progression to a downstream transpososome state.

The cryo-EM map of TnsC_3_–TnsD_1_–DNA_1_ complex allowed model building of the N-terminal domain (NTD) of TnsD, comprising two helix-turn-helix domains (HTH1 and HTH2) connected by a zinc finger domain (ZnF1), as well as a second zinc finger (ZnF2) and HTH3 from the C-terminal domain (CTD) (**Fig. 2a,c**). Three TnsC protomers in the AMP-PNP-bound state were also modeled (**Extended Data Fig. 5a**), with TnsC.1 and TnsC.2 modeled from 6–395, and TnsC.3 modeled from 7–423. TnsC.3 contains a self-inhibitory structure (residues 396–423) as described in Tn7 system^34^. We built a DNA model encompassing 47 bp of the *glmS* gene (positions −47 to −1) and 11 bp of downstream DNA (**Fig. 1c,f and Supplementary Fig. 3a**).

**Fig. 2.**
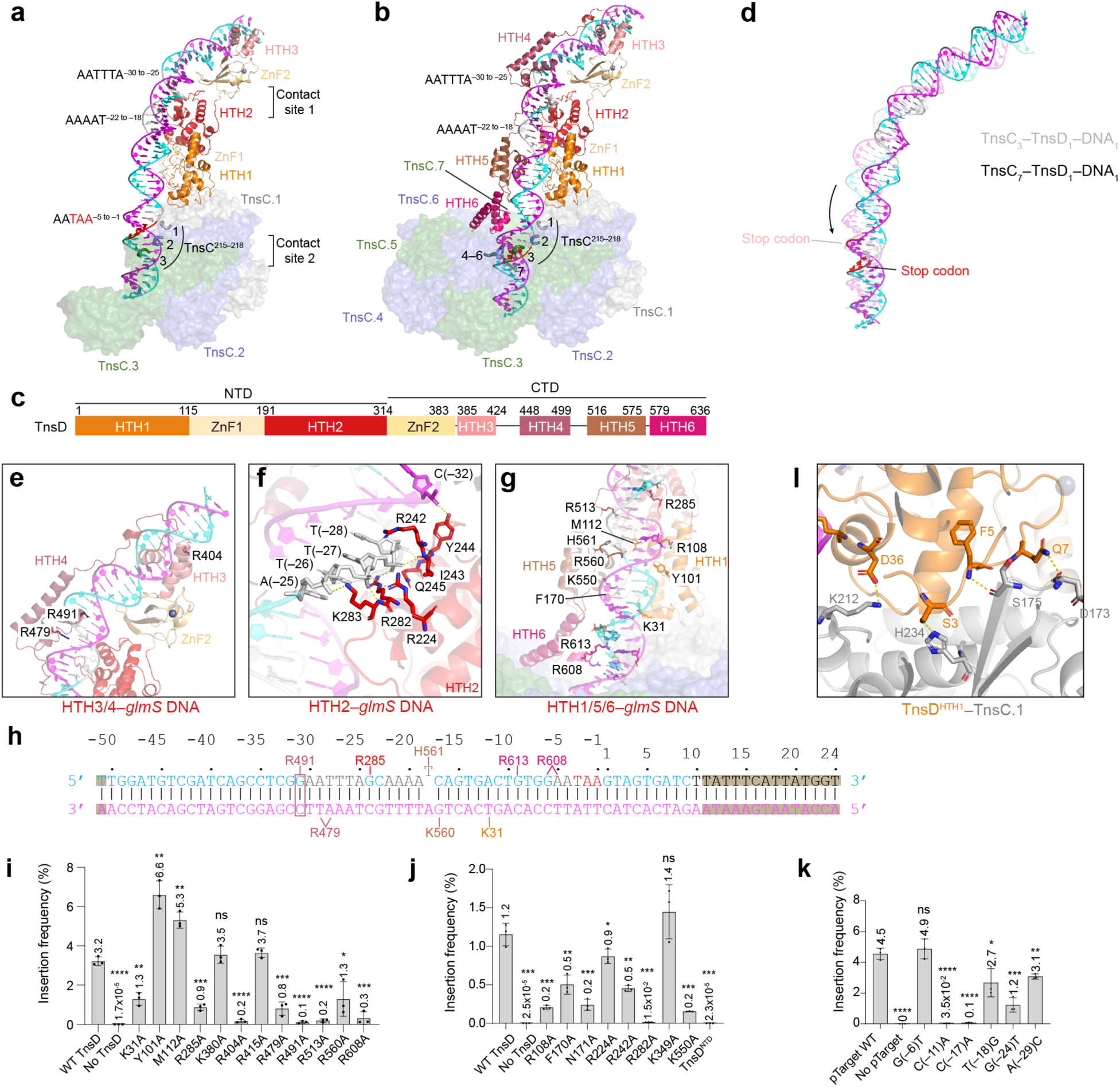
TnsD recognizes the *glmS* attachment site in two steps. **a,b**, Structures of the TnsC_3_–TnsD_1_–DNA_1_ early recognition state (**a**) and the TnsC_7_–TnsD_1_–DNA_1_ late recognition state (**b**), showing progressive engagement of the *glmS* target DNA by TnsD domains and TnsC. A/T-rich DNA segments are indicated. **c**, Domain organization of *Av*TnsD. **d**, Superimposition of target DNA from the TnsC_3_–TnsD_1_–DNA_1_ and TnsC_7_–TnsD_1_–DNA_1_ structures, showing DNA remodeling during TnsC assembly. **e–g**, Close-up views of TnsD–*glmS* DNA interactions mediated by HTH3/HTH4 (**e**), HTH2 (**f**) and HTH1/HTH5/HTH6 (**g**). **h**, Summary schematic of TnsD–*glmS* DNA contacts. Shaded regions indicate nucleotides that were not modeled in the structure. **i,j**, Transposition activities of TnsD mutants. **k**, Transposition activities of *glmS* target mutants. Data are presented as mean values ± SD (n = 3). The stars indicate the significance of the data compared with the wild type. ∗p < 0.05, ∗∗p < 0.01, ∗∗∗p < 0.001, ∗∗∗∗p < 0.0001, ns, not significant. **l**, Interface between TnsD^HTH1^ and TnsC.1 in TnsC_7_-TnsD_1_-DNA_1_.

The TnsC_4_–TnsD_1_–DNA_1_ structure represents a step forward in the assembly process with an additional TnsC.4. As expected from a stepwise assembly process, the self-inhibitory helix observed in the terminal TnsC subunit is released from TnsC.3 and appears in TnsC.4 (**Fig. 1c,d and Extended Data Fig. 5a**). The TnsC_7_–TnsD_1_–DNA_1_ structure contained a complete heptameric TnsC ring and the entire TnsD protein. Because initial reconstructions showed preferred orientation (**Supplementary Fig. 2**), we collected a tilted dataset, yielding an improved 3.23 Å reconstruction (**Fig. 1e, Extended Data Fig. 3d–g, and Extended Data Table 1**). The improved map enabled modeling of the entire TnsD, including the HTH4–HTH6 domains (**Fig. 1e,f**) and assignment of DNA sequence from positions −50 to −1 of *glmS* and 10 bp of downstream DNA (**Supplementary Fig. 3b**). Together, these structures capture a progression from early TnsD-mediated target engagement to a fully assembled TnsC heptamer on remodeled *glmS* DNA.

### TnsD recognizes *glmS* DNA in two steps

*Av*TnsD shares 38% sequence similarity and 23% identity to TnsD in *E. coli* Tn7 (*Eco*TnsD) and contains two extra domains: ZnF2 and HTH3 (**Fig. 2a–c, Supplementary Fig. 4a,b**). Previous studies showed that *Eco*TnsD interacts with around 36 base pairs at 3′ end of the *glmS*^36^.The extra domains interact with the *glmS* DNA spanning from around –45 to –33 (**Fig. 2b,e**) suggesting the extra domains in *Av*TnsD provide extended DNA recognition sites upstream. Transposition assay demonstrated that *Av*TnsD recognizes not only the *glmS* in *Anabaena variabilis*, but also the endogenous *glmS* in *E. coli*^2^, which has 66.7% sequence identity at the 3′ end of the coding region (**Supplementary Fig. 4c**), indicating a conserved recognition mechanism.

The TnsC_3_–TnsD_1_–DNA_1_ and TnsC_7_–TnsD_1_–DNA_1_ structures represent two stages of *glmS* DNA recognition. In the first step, two interaction surfaces are observed between *glmS* DNA and the TnsC and TnsD proteins: (i) nucleotides –29 to –23 interact with TnsD HTH2 domain, and (ii) nucleotides –3 to 7 contact TnsC protomers (**Fig. 2a and Extended Data Fig. 6a**). At the first interface, the DNA phosphate backbone is stabilized by a network of hydrogen-bonding interactions with TnsD side chains and main-chain amide groups (**Fig. 2f**). At the second, the DNA fragment interacts with TnsC protomers, with the loop TnsC^215–218^ positioned along the major groove (**Fig. 2a**). The interactions at these two regions induce bending of *glmS* DNA into an ‘arc-like’ structure (**Fig. 2a**). ZnF2 and HTH3 also contact DNA; however, limited resolution precludes identification of detailed interactions (**Extended Data Fig. 3b**).

Sequence-specific recognition is primarily mediated by HTH4–6 as observed in the TnsC_7_–TnsD_1_–DNA_1_ structure **(Fig. 2b)**. Superimposition of TnsC_3_–TnsD_1_–DNA_1_ and TnsC_7_–TnsD_1_–DNA_1_ shows that ZnF2 and HTH3 domains rotated with the *glmS* DNA (**Fig. 2d and Supplementary Video 1**). Each HTH domain contributes to target recognition by base recognition through hydrogen bonds and backbone stabilization through electrostatic interactions (**Fig. 2g and Extended Data Fig. 6b,d**). Specifically, in HTH4, R479 interacts with T(–29) and A(–28), while R491 interacts with base pairs G(–31) and C(–31) (**Fig. 2e,h and Extended Data Fig. 6b,d**). In HTH5, K560 forms a hydrogen bond with G(–17). In HTH6, R608 forms a hydrogen bond with G(–6) and A(–5), and R613 forms hydrogen bonds with G(–9) and A(–8) (**Fig. 2g and Extended Data Fig. 6b,d**). Consistent with the structural model, mutations of key recognition residues significantly reduced the transposition activity (**Fig. 2i,j and Extended Data Fig. 6c**). Correspondingly, mutation of key nucleotides in the target DNA (e.g., G(–17)) also markedly reduces transposition activity (**Fig. 2k**).

The interaction network between HTH2 and *glmS* DNA is maintained in the TnsC_7_–TnsD_1_–DNA_1_ structure (**Fig. 2f**). Mutating the key residues R224, R242, and R282 to alanine significantly reduced their activity, indicating the importance of these interactions (**Fig. 2j**). Segment from –30 to –25 is A-T rich and undergoes the most severe DNA bending, which suggests local DNA flexibility is another important factor for target recognition^24,40,41^.

More hydrogen bonds were established between HTH1 and *glmS* DNA as the DNA moved closer to the NTD (**Extended Data Fig. 6a,b**). For example, K31 in HTH1 used to be far away from the *glmS* DNA but forms hydrogen bonds with the bases of G(–11) and T(–12) (**Fig. 2g**). One unexpected observation is that M112 from HTH1 domain inserts between the DNA bases and causes the kink at T(–18), with the flipped T(–18) stabilized by the H561 by a hydrogen bond (**Extended Data Fig. 6b,d**). However, mutation of M112 to alanine surprisingly increased the insertion frequency from 3.2% to 5.3% (**Fig. 2i**). Similar result was found when mutating Y101 to alanine (**Fig. 2i**). The bulky side chain of tyrosine and methionine may impose steric constraints that hinder DNA conformational changes during target recognition. Sequence alignment shows that both Y101 and M112 in *Av*TnsD corresponds to an alanine residue in *Eco*TnsD (**Supplementary Fig. 4b**), which supports this interpretation. Besides, residue K550 from HTH5 stacks with F170 from HTH1 (**Extended Data Fig. 6e**) and mutating any one of them to alanine reduced the transposition significantly, suggesting that intramolecular communication within TnsD contributes to target recognition (**Fig. 2j**). Together, these data indicate that the DNA conformational change arises from coordinated interactions between HTH domains and *glmS* DNA.

### Stepwise TnsC oligomerization remodels the target DNA

Recognition of *glmS* DNA by TnsD appears to be coordinated with the stepwise TnsC oligomerization. The TnsC protomer that interacts with TnsD (TnsC.1) is recruited to the target site through direct contacts with the HTH1 and ZnF1 domains of TnsD (**Fig. 2l**). L108–T139 in TnsC.1, which forms the interaction surface for TnsC.1 and TnsD interaction, is in a different conformation in the remaining TnsC protomers (**Extended Data Fig. 5a**). Consistent with previous observations for *Eco*TnsC, L108–T139 from TnsC.2 contacts the nearby TnsC.1 at the interface between the large and small AAA+ domains (**Extended Data Fig. 5b**)^34,35^. At the catalytic site of TnsC.1, the arginine finger R292 and R293 from TnsC.2 interact with the phosphate group of AMP-PNP bound to TnsC.1 (**Extended Data Fig. 5c**), thereby stabilizing the recruitment of the second TnsC subunit. TnsC.3 engages TnsC.2 through similar interactions, but an additional helix from I411 to D423 in TnsC.3, together with the upstream loop from D396 to D410, sterically occludes interactions between the arginine finger of TnsC.4 and the AMP-PNP bound to TnsC.3, thereby preventing the recruitment of a fourth TnsC subunit (**Extended Data Fig. 5d**).

In the TnsC_4_–TnsD_1_–DNA_1_ structure, the fourth TnsC is loaded to the TnsC trimer, where the steric obstacle in TnsC.3 is released. Addition of TnsC.4 happened together with the local conformational changes of *glmS* DNA moving towards the TnsC.4 (**Extended Data Fig. 5e**). Similar but more obvious conformational changes were observed when comparing the TnsC_3_–TnsD_1_–DNA_1_ and TnsC_7_–TnsD_1_–DNA_1_ (**Extended Data Fig. 5f**), indicating accumulative conformational changes of *glmS* DNA and TnsD as TnsC.4–7 subunits are loaded to form the TnsC heptamer. Interestingly, in the TnsC_7_–TnsD_1_–DNA_1_ structure, the *glmS* DNA is shifted downward by approximately 4 bp (**Fig. 2d**). As a result, although the same TnsC residues mediate DNA binding in both assemblies, these interactions occur predominantly along the DNA minor groove in the heptameric complex, in contrast to major-groove interaction observed in the trimer complex (**Fig. 2a,b and Extended Data Fig. 5g,h**).

### ATP supports progression to an asymmetric STC

In the presence of ATP, the assembly progressed to an asymmetric STC, which was resolved at 3.19 Å resolution (**Fig. 3a–c, Extended Data Fig. 2f–h, and 4, and Extended Data Table 1**). In contrast to previously reported STC structures^20,21,24^, which contain a single TnsB-binding site at each transposon end, the TnsB-DNA assemblies at the RE and LE in our structure differ, with a more ordered RE spanning three TnsB binding sites, whereas the LE contains only one (**Fig. 3d,e**). Overall, the structure contains two TnsA bound to TnsC C-terminal tails, five copies of TnsB, RE and LE of the transposon built to 54 and 42 bp, respectively, and the target DNA of 45 bp (**Fig. 3a–d**).

**Fig. 3.**
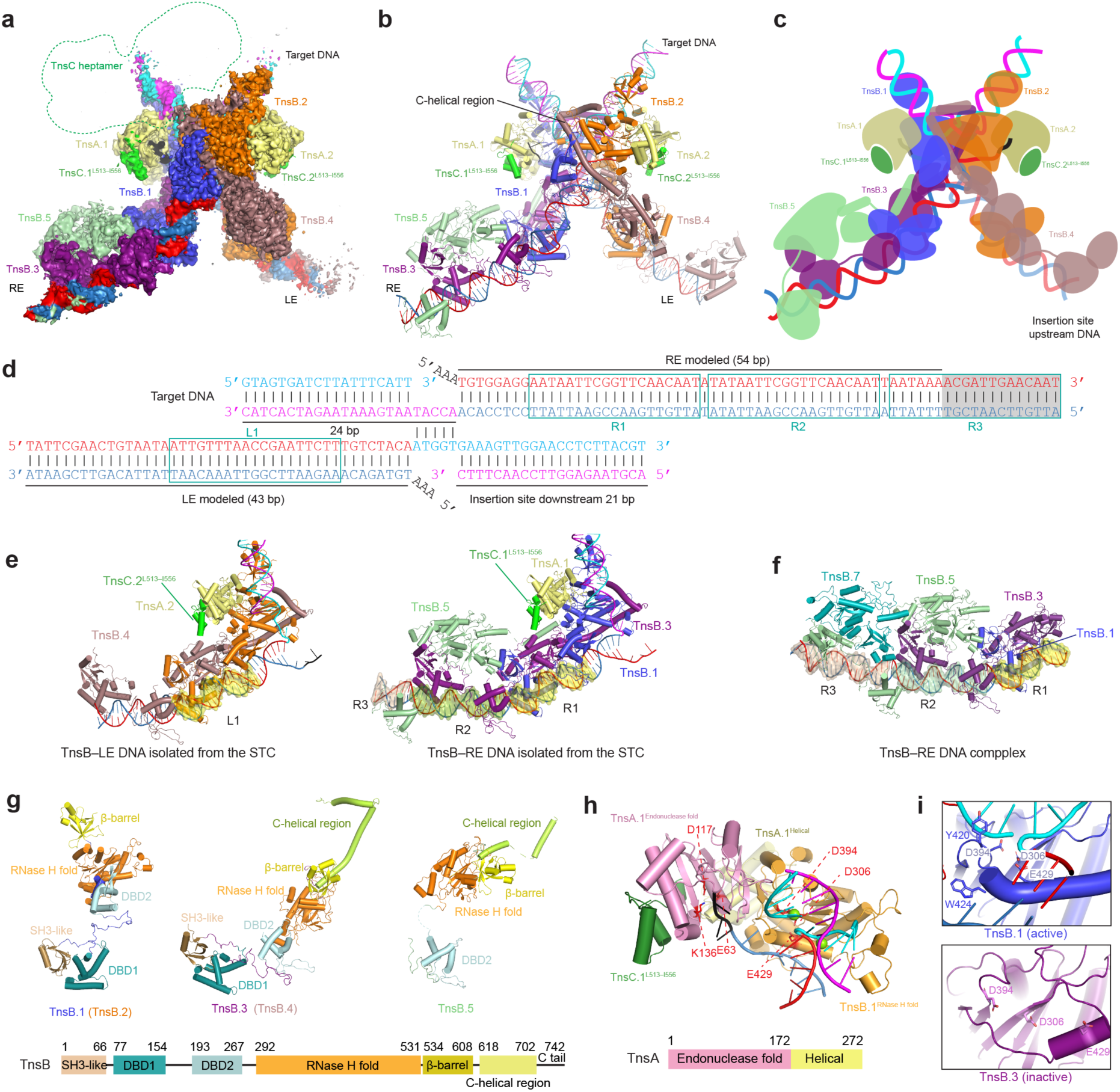
An asymmetric STC defines insertion orientation in *Av*CAST. **a**, Cryo-EM density map of the *Av*CAST STC assembled in the presence of ATP. The modeled position of the TnsC heptamer, inferred from the target-bound TnsC–TnsD–DNA assembly, is indicated by a dashed outline; no corresponding density was resolved in the STC map. **b**, Atomic model of the STC shown in the same orientation as in **a**. **c**, Schematic of the asymmetric STC architecture. **d**, Schematic of the DNA arrangement in the STC, showing the target DNA sequence, insertion site, modeled LE and RE regions, and TnsB-binding sites. Shaded regions indicate nucleotides that were not modeled in the structure. **e**, TnsB–LE and TnsB–RE modules extracted from the STC, showing asymmetric engagement of the two transposon ends. **f**, Structure of the isolated TnsB–RE DNA complex, shown for comparison with the RE-bound module in the STC. **g**, Domain organization and conformations of TnsB protomers in the STC. **h**, TnsA–TnsB interface at the active transposase site. **i**, Comparison of active and inactive TnsB active-site configurations.

Consistent with our previous model of the type I-B2 system^24^, four TnsB molecules and two TnsA molecules form the core of the STC **(Fig. 3a–c)**, whereas type V-K CASTs lack TnsA yet retain a similar overall architecture^20,21^. TnsB subunits proximal to the insertion site (TnsB.1 and TnsB.2) act as active nucleases that catalyze the reaction in trans: TnsB.1 recognizes the RE via its DBD1 and DBD2 domains but cleaves and rejoins the LE at the target site, whereas TnsB.2 reciprocally recognizes the LE and catalyzes the RE (**Fig. 3b,e,g**). TnsA and TnsB interact through β-strand extension, as observed in *Pmc*CAST^24^, positioning their catalytic sites at the target site loaded with substrate DNA (**Fig. 3h**). The TnsC C-terminal region (TnsC^513–556^) laterally contacts and recruits TnsA through predominantly hydrophobic interactions, resembling the TnsA–TnsC^504–555^ crystal complex in Tn7 (**Supplementary Fig. 5a,b**). In contrast, in *Pmc*CAST, the TnsC C-terminal region is positioned at the junction between the TnsA and TnsB interface^24^. This difference likely reflects that TnsA and TnsB in *Av*CAST are separate proteins requiring TnsC-mediated recruitment of TnsA, whereas in *Pmc*CAST such recruitment is unnecessary due to the TnsAB fusion.

Distal TnsB subunits (TnsB.3 and TnsB.4) primarily serve structural roles. TnsB.4 (or TnsB.3) stabilizes the complex: its C-helical region interacts with both TnsB.1 and TnsB.2 and reinforces the catalytic domain of TnsB.2 (or TnsB.1) at the insertion site (**Fig. 3b,c,g**). In these subunits, the catalytic residue E429 is displaced from D394 and D306, rendering them inactive (**Fig. 3i**). Superimposition of the catalytic domain of TnsB.1 and TnsB.3 shows a conformational change at the helix harboring E429. Y420 and W424 in TnsB.1 interact with target DNA and transposon end, respectively, driving repositioning of E429 into the active configuration. Activation of TnsB nuclease requires transpososome assembly, which appears to be a conserved mechanism in *Sh*CAST and prototypical Tn7^21,33^ (**Fig. 3i**).

TnsB.5 is resolved with the DBD2, catalytic domain, β-barrel, and a partial C-helical region modeled **(Fig. 3g)**. An α-helix spanning V644 to Y666 within the C-helical region of TnsB.5 engages the catalytic domain of TnsB.3, similar to that between TnsB.2–TnsB.4 interface (**Fig. 3b**). The RE-bound TnsB subunits (TnsB.1, TnsB.3, and TnsB.5) are positioned closer to the expected TnsC heptameric ring, which may contribute to the orientation of transposon insertion (**Fig. 3a**).

### TnsB domains in the STC core are flexible in the TnsB–RE DNA complex

To define the TnsB–DNA state before STC formation, we assembled a TnsB–DNA complex by incubating purified TnsB and 80 bp RE DNA with three TnsB-binding sites (**Extended Data Fig. 2a**) and determined its structure at 3.9 Å resolution (**Extended Data Fig. 7a–f and Extended Data Table 1**).

Using the TnsB–DNA model distal to the insertion site, we built a TnsB–DNA model containing four TnsB molecules bound to the RE DNA (**Fig. 3f and Extended Data Fig. 7g**). The DBD1 and DBD2 domains of TnsB.1, TnsB.3, TnsB.5 bind the R1, R2, and R3 sites, respectively, while a partial TnsB.7 contacts the DBD1 domain of TnsB.5 (**Extended Data Fig. 7h**). The C-terminal helical domains of TnsB that mediate inter-subunit interactions in STC are not resolved in the TnsB–DNA structure. Notably, only DBD1 of TnsB.1 is visible in the density map, suggesting that the remaining domains of TnsB.1 are flexible. These observations support an STC assembly model in which the flexible TnsB.1 domains, together with TnsA, TnsC^L513–I556^, and the C-terminal helix of TnsB.3, engage the opposite transposon end, interacting with TnsB.2 and the target DNA to enable catalysis (**Fig. 3e,f**).

### Target recognition by *Av*Cascade and TniQ recruitment

To understand the mechanism of the RNA-guided pathway, we assembled the *Av*Cascade–TniQ–DNA complex using a pull-down assay (**Extended Data Fig. 8a**). The DNA substrate contained an 11-bp upstream sequence with an “ATG” PAM, a 36-bp protospacer, and 43-bp downstream sequence. Cryo-EM analysis resolved three distinct structures: two *Av*Cascade–DNA complexes, *Av*Cascade–DNA-1 (2.64 Å; 63.3%) and *Av*Cascade–DNA-2 (3.01 Å; 19.7%), exhibiting stabilized density near the PAM-proximal or PAM-distal region, and an *Av*Cascade–TniQ–DNA (3.16 Å; 17.0%) in which TniQ binds at the PAM-distal region (**Fig. 4a-d,h, Extended Data Fig. 8b–h, and Extended Data Table 1**). *Av*Cascade is composed of Cas5, Cas6, Cas7, Cas8, Cas11, and crRNA with a stoichiometry of 1:1:7:1:3:1. Similar to *Pmc*Cascade, Cas11 is internally translated from the *cas8* gene.

**Fig. 4.**
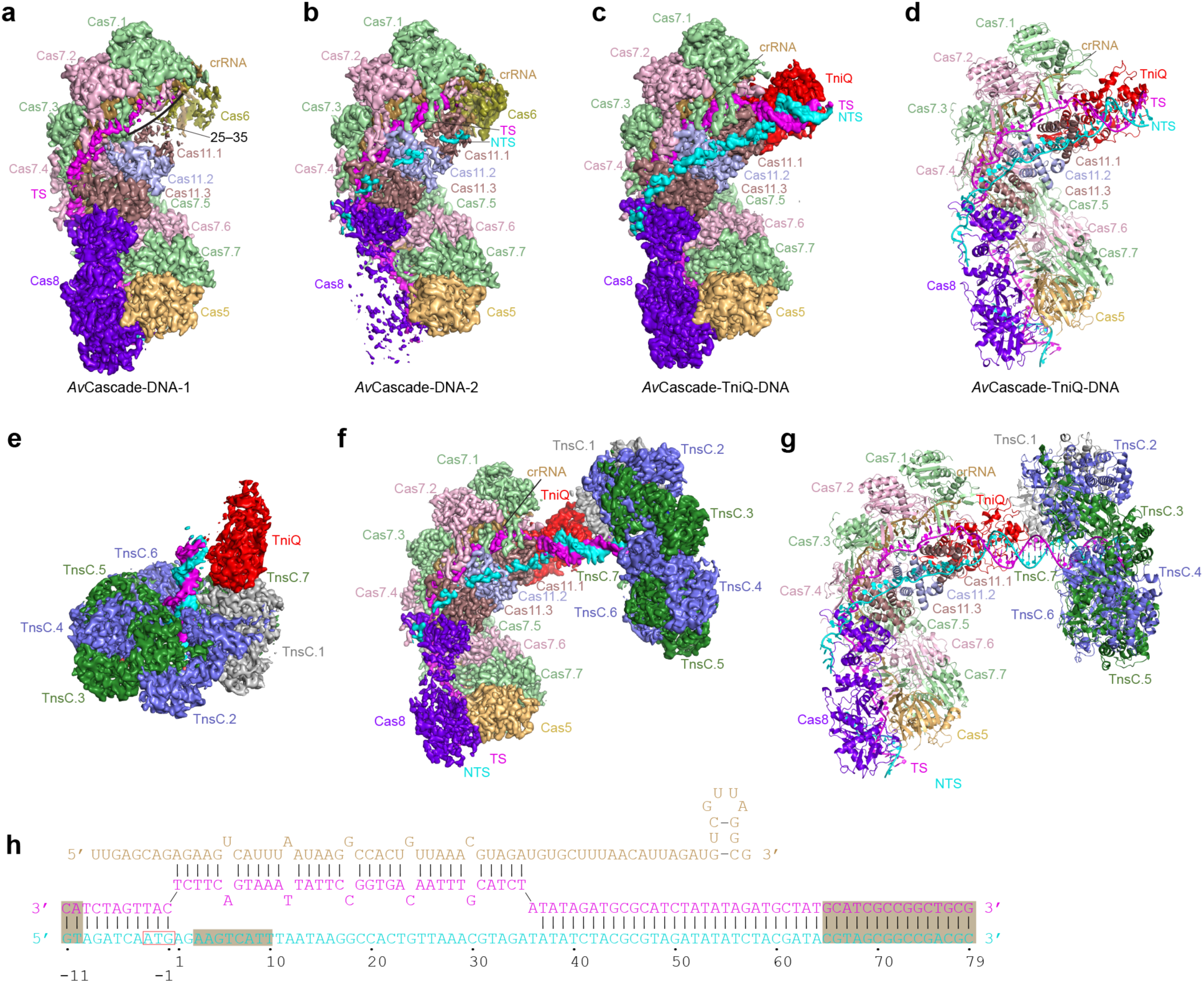
RNA-guided targeting recruits TniQ and positions TnsC on target DNA. **a,b**, Cryo-EM maps of *Av*Cascade–DNA assemblies. Cascade subunits, crRNA, target strand (TS) and non-target strand (NTS) are indicated. **c,d**, Cryo-EM map (**c**) and atomic model (**d**) of the *Av*Cascade–TniQ–DNA complex, showing recruitment of TniQ to the Cascade-bound target. **e**, Cryo-EM map of the TniQ–TnsC–DNA assembly, showing TniQ associated with a TnsC heptamer on DNA. **f,g**, Composite cryo-EM map (**f**) and model (**g**) of the RNA-guided targeting assembly, generated by superimposing the TniQ–DNA module from the *Av*Cascade–TniQ–DNA and TniQ–TnsC–DNA structures. **h**, Schematic of the crRNA–target DNA pairing and PAM region recognized by *Av*Cascade. Shaded regions indicate nucleotides that were not modeled in the structure.

The three structures indicate a sequential target-recognition process for TniQ recruitment. *Av*Cascade–DNA-1 exhibits the highest resolution at the PAM-proximal region but weaker density at positions 25–35 of the RNA–DNA heteroduplex (**Fig. 4a**), suggesting that it represents an earlier stage of target recognition. Cas5 and Cas8 recognize the “ATG” PAM, consistent with *Pmc*CAST ^23^ (**Fig. 5a and Extended Data Fig. 8i,j**), rather than the previously reported “AT” PAM^2^. Consistently, the target plasmid containing an “ATG” PAM displayed higher transposition activity than the plasmid containing an “AT” PAM **(Extended Data Fig. 1o)**. R75 and K77 in Cas5 play critical roles in PAM recognition by interacting with the backbone of the target strand (**Fig. 5b,c**). Alanine substitution of either residue significantly reduced RNA-guided transposition activity (**Fig. 5d and Extended Data Fig. 8k**). Q296, H117, and K119 in Cas8 interact with the bases C(−1), A(−2), and T(−3) of the target strand, respectively.

**Fig. 5.**
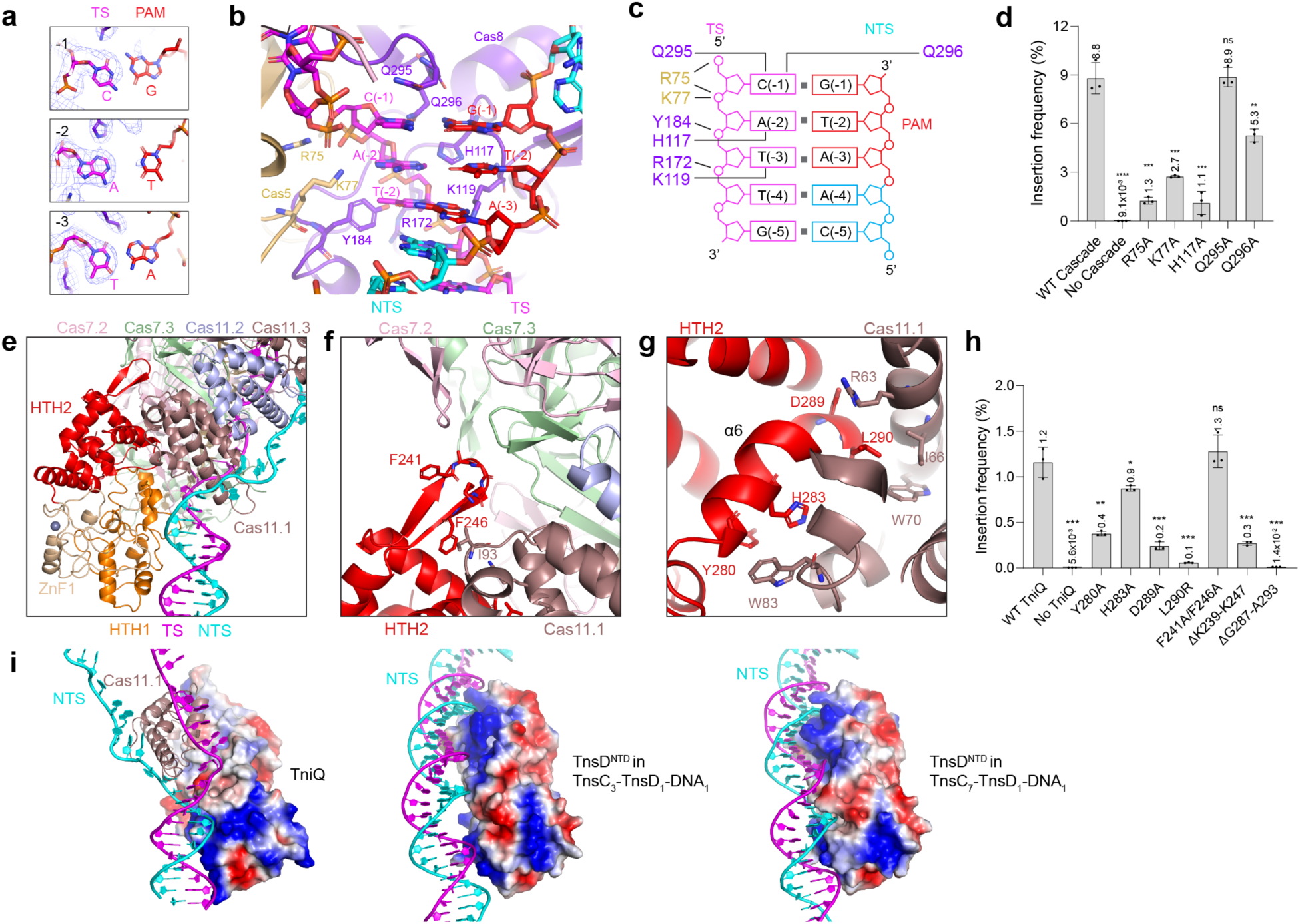
Cascade recognizes an ATG PAM and recruits TniQ. **a**, Cryo-EM density for the PAM nucleotides in the *Av*Cascade–TniQ–DNA structure. **b**, Close-up view of PAM recognition by Cas8 and Cas5. **c**, Schematic of PAM recognition, showing interactions between Cascade residues and the PAM. **d**, Transposition activities of Cascade PAM-recognition mutants. **e**, Close-up view of TniQ in the *Av*Cascade–TniQ–DNA structure. **f,g**, Close-up views of TniQ–Cascade interfaces. **h**, Transposition activities of TniQ mutants. Data are presented as mean values ± SD (n = 3). The stars indicate the significance of the data compared with the wild type. ∗p < 0.05, ∗∗p < 0.01, ∗∗∗p < 0.001, ns, not significant. **i**, Electrostatic surface comparison of TniQ and TnsD^NTD^ in RNA-guided and TnsD-guided targeting assemblies. Analogous DNA-facing surfaces are observed in the *Av*Cascade–TniQ–DNA and TnsC_7_–TnsD_1_–DNA_1_ structures.

After PAM recognition, the crRNA pairs with the target strand. Pairing at positions 25–35 completes R-loop formation and is accompanied by conformational changes in the Cas11 filament (**Supplementary Video 2**), which further stabilize the non-target strand through interactions with three copies of Cas11. Structural superimposition of the *Av*Cascade–TniQ–DNA and *Av*Cascade–DNA complexes reveals an overlap between the position occupied by Cas6 and the bound TniQ (**Fig. 4b,c**), suggesting that TniQ binding displaces Cas6, rendering it flexible.

### TniQ is recruited by interacting with Cas11

*Av*Cascade recruits TniQ mainly through Cas11 (**Fig. 5e**). Specifically, a loop in the antiparallel β-strand in HTH2 domain is positioned at a junction formed by loops from Cas7.2, Cas7.3, and Cas11.1, with hydrophobic contact between I93 in Cas11.1 and F246 in TniQ (**Fig. 5f**). In contrast, the α6 helix forms the primary interface with Cas11.1: Y280 and H283 pack against W83, D289 forms an electrostatic interaction with R63, and L290 engages I66 and W70 through hydrophobic contacts (**Fig. 5g**). Replacement of the loop (ΔK239–K247) or part of α6 helix (ΔG287–A293) with a “GSA” linker impaired activity, highlighting the importance of these interaction surfaces (**Fig. 5h and Extended Data Fig. 8l**). Consistent with these interactions, mutations Y280A, H283A, D289A, and L290R each significantly reduced RNA-guided transposition activity (**Fig. 5h**).

### TniQ binds DNA in a state ready for TnsC heptamerization

Our structures show that the HTH1 domain of TniQ is already engaged with DNA at a position comparable to that observed in the TnsC_7_–TnsD_1_–DNA_1_ complex (**Fig. 5i**), suggesting that this configuration primes the DNA for TnsC heptamer assembly. To test this, we determined a cryo-EM structure of the TniQ–TnsC–DNA complex (**Fig. 4e, Extended Data Fig. 9, and Extended Data Table 1**), which adopts a stoichiometry of 1:7:1, with no evidence for a stable 1:3:1 intermediate. We further generated a model of the *Av*Cascade–TniQ–TnsC–DNA assembly by superposing TniQ and DNA from both structures (**Fig. 4f,g**). The model indicates that, following *Av*Cascade-mediated target recognition, TniQ is poised to engage TnsC, positioning TnsC.1 at the DNA minor groove (**Fig. 4g and Fig. 5i**).

Finally, to generate a structural model for insertion site selection, we built a model of the TnsABCD transpososome by aligning the TnsC_7_–TnsD_1_–DNA_1_ and STC structures to the TnsABCD transpososome structure from *Pmc*CAST ^24^. Notably, the DNA models align well, placing the insertion site 24 bp downstream of the *glmS* stop codon, consistent with the insertion pattern observed for TnsD-guided transposition (**Extended Data Fig. 1f,j and 10a**). Similarly, we generated a complete transpososome model in the RNA-guided pathway by aligning *Av*Cascade-TniQ–TnsC–DNA and STC structures, resulting in an insertion site 79 bp downstream of the “ATG” PAM, consistent with the predominant insertion site observed *in vitro* (**Extended Data Fig. 1l–n, and 10b**). Modeling of the complete transpososome reveals a steric clash between the SH3-like domain of TnsB.1 and the TnsC heptamer (TnsC.4), which may explain why a stable TnsC heptamer cannot coexist with the STC (**Extended Data Fig. 10c**). Together, these results establish a structural framework for understanding the TnsD- and RNA-guided transposition pathways in type I-B1 CASTs.

## Discussion

Here, we define how target recognition is converted into productive and directional DNA insertion in *Av*CAST (**Fig. 6**), a type I-B1 CRISPR-associated transposon closely related to prototypical Tn7. By comparing protein-guided and RNA-guided transposition, we show that DNA remodeling acts as a key signal that couples target recognition to transpososome assembly, converging on a shared architecture in which TnsC is positioned along the DNA minor groove. Together, these findings identify DNA remodeling and TnsC positioning as key features that underlie productive transpososome assembly.

**Fig. 6.**
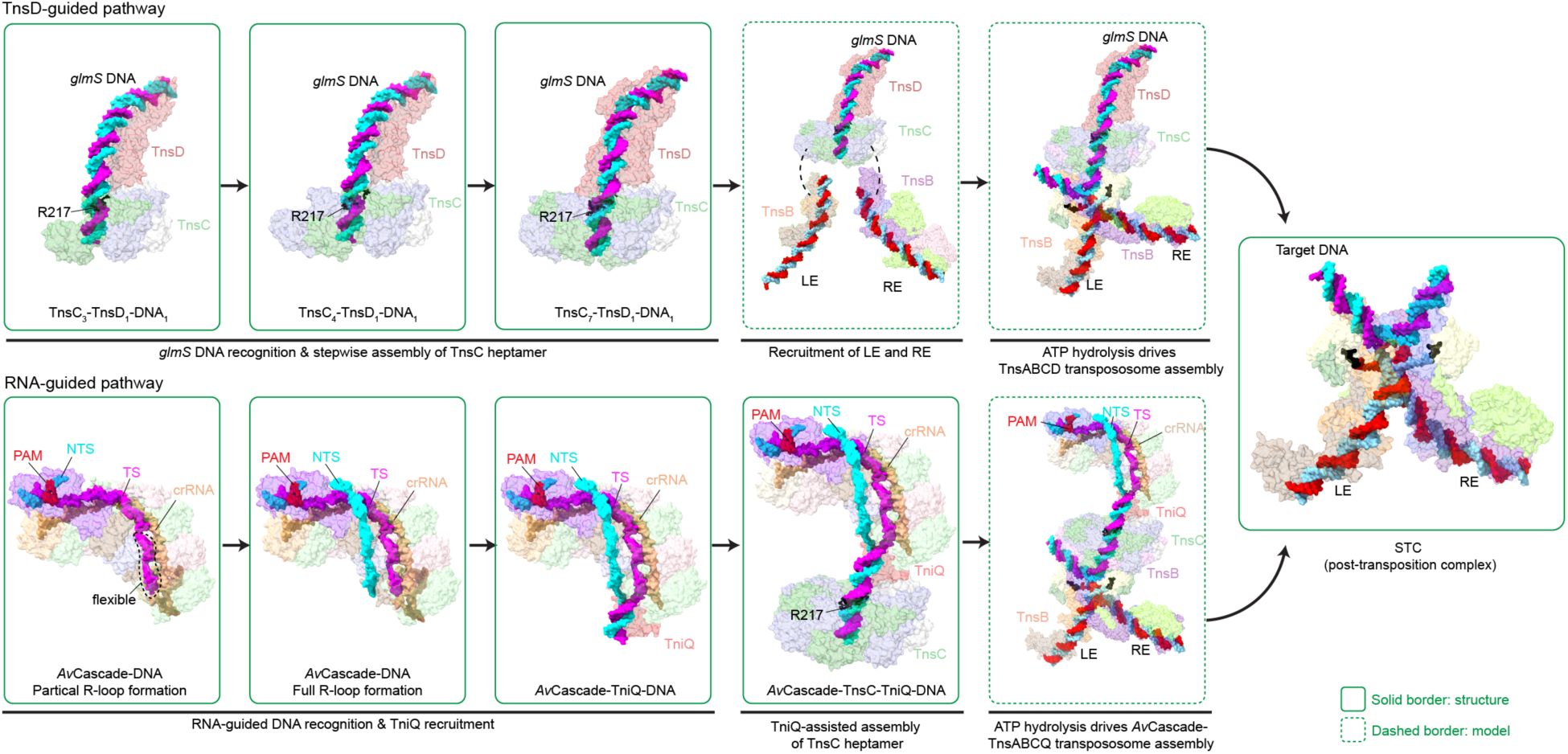
Model for convergence of protein- and RNA-guided targeting in *Av*CAST. In the TnsD-guided pathway, TnsD recognizes the *glmS* attachment site and remodels the target DNA during stepwise TnsC assembly. Progressive loading of TnsC converts the target into a minor-groove-associated TnsC heptameric state that is competent for recruitment of LE and RE and downstream TnsA/TnsB transpososome assembly. In the RNA-guided pathway, *Av*Cascade recognizes the target through PAM-dependent R-loop formation and recruits TniQ. TniQ positions TnsC on target DNA in a geometry analogous to the TnsD-guided pathway, thereby linking CRISPR-guided recognition to the shared TnsABC machinery. ATP supports progression from target-bound TnsC assemblies to an asymmetric STC, coupling target recognition to directional DNA insertion.

While sharing features with prototypical Tn7, *Av*CAST exhibits key mechanistic differences in how target recognition is coupled to TnsC loading. A study of truncated *Eco*TnsD from Tn7, encompassing only the NTD, showed that *Eco*TnsD^NTD^ functions as a nucleotide exchange factor, facilitating TnsC loading at the target site by promoting ADP-ATP exchange within the TnsC dimer^34^. In contrast, *Av*TnsC is purified as a monomer lacking bound nucleotide (**Extended Data Fig. 5i,j**), suggesting that *Av*TnsD^NTD^ may not rely on a nucleotide exchange mechanism. The differences underscore mechanistic diversity even among closely related systems.

Our findings provide insight into the long-standing question of how insertion orientation is established in Tn7 and CAST systems. Although asymmetric TnsB-binding sites at the two transposon ends have been proposed to contribute to directional insertion^1^, structural studies of STCs or transpososome complexes in V-K^20–22,42^, I-B2^24^, and I-F^43^ CAST systems reveal largely symmetric architectures, with only a single TnsB-binding site resolved at each end. In contrast, the *Av*CAST STC exhibits a distinct asymmetry, with RE containing three contiguous TnsB-binding sites and forming a more stable and well-ordered transposon-end complex positioned toward the TnsC heptamer (**Fig. 3a,e**). This organization suggests that the more stable RE may be preferentially positioned near the TnsC heptamer controlling the insertion orientation of the paired-end complex during transpososome formation. Alternatively, the two ends may be recruited sequentially, with the more stable RE engaging first, followed by incorporation of the less stable LE. Although we do not observe extensive direct contacts between RE-associated TnsB subunits and TnsC, transient interactions during assembly may be lost upon formation of the final STC. Together, these observations suggest that insertion orientation may arise from asymmetric transposon-end organization established during assembly. Further structural and biochemical studies, particularly of assembly intermediates, will be required to define how these features contribute to directional insertion.

Although ATP hydrolysis has been implicated in target-site selection in some Tn7-like systems, including Tn7 and type V-K CASTs^17,44^, where impaired hydrolysis can increase off-target insertion, our data show that AMP-PNP still supports accurate TnsD-guided target recognition in *Av*CAST **(Extended Data Fig. 1d,g)**. The residual activity observed with AMP-PNP suggests that a small fraction of target-bound assemblies can occasionally progress to productive transpososomes, although ATP hydrolysis greatly increases the efficiency of this transition. Thus, ATP-dependent nucleotide turnover appears to act primarily after target recognition in this system, promoting progression from target-bound assemblies to the STC. Together with the conserved TnsC–TnsB C-terminal-hook interaction observed in Tn7-like transposons^20,24,45^, our STC structure supports a model in which ATP-dependent remodeling of the TnsC assembly could be transmitted through the TnsB C-terminal hook/region to promote assembly of a catalytically competent transpososome. This model is consistent with observations in IS21-family transposons, in which ATPase-driven nucleotide turnover before strand transfer regulates transposase assembly and conformational changes that position catalytic residues for strand transfer^46^.

Our results also provide guidance for engineering CAST-based genome-editing systems. First, our structural and biochemical analyses refine the functional PAM requirement of *Av*CAST to ATG, which will inform retargeting of this system. Second, our structures show that productive transposition requires not only recruitment of TnsC to target DNA but also proper positioning of the TnsC oligomer along the DNA minor groove. This principle may help explain why simple programmable DNA-binding module–TniQ fusions do not always support efficient targeted transposition^47^, and suggests that engineering efforts should consider not only guide-dependent recruitment of CAST components but also the spatial organization required for productive transpososome assembly. Consistent with the potential of structure-guided engineering, recent computational protein-design work generated a simplified *Pmc*CAST-derived system with improved activity^48^.

In summary, our study defines how type I-B1 CASTs integrate distinct targeting pathways into a shared assembly program, establishing a framework for understanding CAST function and guiding the engineering of programmable gene insertion systems with improved specificity, efficiency, and control over insertion orientation.

## Methods

### Plasmid construction

The primers used in this study are listed in **Supplementary Table 1**. Three plasmids, pCDFDuet_Cas6_CRISPR array, pRSFDuet_Cas7_Cas5, pETDuet_Cas8, were constructed for expression of *Av*Cascade in *E. coli* BL21(DE3). Genes encoding the Cascade components were amplified by PCR from pHelper(*Av*CAST)_*Av*PSP1 (Addgene #168133) and inserted into the corresponding Duet vectors using the In-Fusion cloning kit (Takara). An N-terminal 6×His tag was fused to Cas8 to facilitate purification of the *Av*Cascade complex.

The gene of *Av*TniQ was amplified from pHelper(*Av*CAST)_*Av*PSP1 and cloned into ColE1-based pTwinStrep-SUMO vector for expression and purification. Point mutations in Cascade subunits, TniQ, TnsD and pTarget were generated using QuickChange site-directed mutagenesis. The loop (ΔK239–K247) and part of the α6 helix (ΔG287–A293) in TniQ were replaced with GSA linkers of matching length, “GSAGSAGS” and “GSAGSAG,” respectively, using an In-Fusion cloning kit.

The pDonor plasmid carrying a kanamycin resistance cassette was constructed by replacing the left and right ends (LE and RE) of a previously reported type V-K donor plasmid (Addgene #127924) with the corresponding RE and LE sequences from the type I-B1 CAST system. Truncations at both transposon ends in the pDonor plasmid were generated by PCR-mediated deletion, followed by self-ligation using the In-Fusion cloning kit.

The target plasmid containing target sites for both RNA-guided and TnsD-guided transposition was constructed by modifying the pTarget plasmid (Addgene #168163) previously used for *Pmc*CAST systems. Specifically, the “ATG” protospacer-adjacent motif (PAM) in the *Pmc*CAST target was converted to “AT” by deletion of the terminal “G”. Notably, the first base of the spacer is “G,” which, based on cryo–EM analysis, contributes to the PAM sequence. In addition, the tRNA-Val gene was replaced with the final 50 bp of the *glmS* gene, including the stop codon.

### Protein expression and purification

TnsB, TnsC, and TnsD were expressed and purified as previously described ^2^. Briefly, all proteins were expressed in *E. coli* BL21(DE3)-RIPL cells. Cultures were grown in Terrific Broth (TB) with 100 μg/mL ampicillin to an OD600 of ∼0.6 and induced with 0.5 mM IPTG at 16 °C overnight. Cells were harvested by centrifugation and resuspended in lysis buffer containing 50 mM Tris–HCl (pH 8.0), 500 mM NaCl, 10% (v/v) glycerol, 1 mM DTT, and 1 mM PMSF. Cells were lysed by sonication, and the clarified lysate was applied to Strep-Tactin Superflow Plus resin (Qiagen). After washing with lysis buffer, proteins were eluted using lysis buffer supplemented with 2.5 mM desthiobiotin (Sigma). Eluted SUMO-tagged proteins were digested with homemade Ulp1-R3 SUMO protease ^49^ (Addgene #113671) at a 1:100 protease-to-substrate molecular ratio at 4 °C for 16 h. The digested samples were concentrated to ∼1 mL and further purified by size-exclusion chromatography using a Superdex 200 Increase 10/300 GL column (Cytiva) equilibrated in 50 mM Tris–HCl (pH 7.5), 500 mM NaCl, 5% (v/v) glycerol, and 2 mM DTT. Peak fractions were pooled, concentrated, flash-frozen, and stored at –80 °C.

TnsA and TniQ were purified using the same strategy as TnsB, TnsC, and TnsD, except that the lysis buffer contained 50 mM Tris–HCl (pH 8.0), 500 mM NaCl, 10% (v/v) glycerol, 1.43 mM β-mercaptoethanol, and 1 mM PMSF. Following SUMO protease digestion, samples were incubated with Ni-NTA resin to remove the cleaved tag prior to concentration and size-exclusion chromatography on a Superdex 75 Increase 10/300 GL column (Cytiva). For TnsA, the column was equilibrated in 50 mM Tris–HCl (pH 7.5), 500 mM NaCl, 5% (v/v) glycerol, and 2 mM DTT, whereas TniQ was purified using 50 mM Tris–HCl (pH 8.0), 500 mM NaCl, 5% (v/v) glycerol, and 1 mM DTT. Final peak fractions were pooled, concentrated, flash-frozen, and stored at –80 °C.

To express the *Av*Cascade complex, plasmids pCDFDuet_Cas6_CRISPR array, pRSFDuet_Cas7_Cas5, and pETDuet_Cas8 were co-transformed into *E. coli* BL21(DE3) cells and selected on LB agar containing 100 μg/mL ampicillin, 50 μg/mL kanamycin, and 50 μg/mL spectinomycin. A single colony was picked and cultured in 50 mL LB broth overnight at 37 °C with antibiotics. Cells were inoculated into TB broth with antibiotics to an OD600 of ∼0.8 and induced with 0.5 mM IPTG at 16 °C overnight. Cells were harvested by centrifugation and resuspended in lysis buffer containing 50 mM Tris–HCl (pH 7.5), 500 mM NaCl, 5% (v/v) glycerol, 1.43 mM β-mercaptoethanol, and 10 mM imidazole, supplemented with Pierce Protease Inhibitor Tablets (Fisher Scientific). Cells were lysed by sonication, and the lysate was clarified by centrifugation at 18,000 rpm for 30 min at 4 °C. The supernatant was filtered and applied to Ni–NTA resin (Qiagen). After washing with lysis buffer supplemented with 30 mM imidazole, bound proteins were eluted with lysis buffer containing 330 mM imidazole. The eluted sample was diluted with buffer containing 50 mM Tris–HCl (pH 7.5), 150 mM NaCl, 5% (v/v) glycerol, and 1 mM DTT, and then loaded onto a HiTrap Q HP column (Cytiva). Proteins were eluted using a step gradient of 0.1–1.0 M NaCl in 50 mM Tris–HCl (pH 7.5), 5% (v/v) glycerol, and 1 mM DTT. Fractions containing the *Av*Cascade complex were pooled, concentrated, and further purified by size-exclusion chromatography on a Superdex 200 Increase 10/300 GL column (Cytiva) equilibrated in 20 mM HEPES (pH 7.5), 200 mM NaCl, 5% (v/v) glycerol, and 1 mM DTT. Peak fractions were pooled, concentrated, flash-frozen, and stored at –80 °C.

### Mass spectrometry analysis and liquid chromatography

Dried peptides obtained were reconstituted in 3% ACN and 0.1% formic acid (FA) before being analyzed by HPLC-MS/MS using equipped with an Ultimate 3000 HPLC (Thermo Fisher Inc., Waltham, MA) and a PEPSEP analytical column (Brucker Scientific), the LC settings and mass spec settings are as described by Grantz et al. 2024. Briefly, 600 ng of peptides were loaded on a trap column and separated on an analytical column. A 130 min elution gradient was constructed by mixing mobile phase solvent A (0.1% FA in water) with solvent B (80% ACN, 0.1% FA in water). Two percent of solvent B was initially used and increased to 12% at 1.6 min, 25% at 80 min, 35% at 100 min, 45% at 105 min, and 95% at 120 min, at which point the gradient was held for 5 min before reverting to 2% at 125.1 min, and 5% at 130 min. An injection of 0.6 µg of sample peptide was performed for each sample. All the data were acquired using an Orbitrap Fusion lumos (Thermo Fisher Inc., Waltham, MA). The mass spectrometry was operated at a positive mode using a data dependent acquisition mode (DDA). The MS settings were kept the same as Grantz et al. 2024 ^50^. The performance of the instrument was analyzed by Hela digest before and after the run.

The raw data analysis was conducted by MS-Fragger ^51^. The analysis settings were kept as default. Resulting protein and peptide intensities were exported and analyzed using Microsoft excel.

### Mass photometry analysis

Mass photometry measurements were performed using a TwoMP instrument (Refeyn, Oxford, UK) operated with AcquireMP software. High-precision Microscope coverslips (No. 1.5H, 24x50 mm, Deckglaser) were cleaned by three sequential cycles of isopropanol and nanopure water, followed by air drying. CultureWall gaskets (3 mm diameter, 1 mm depth; Grace Bio-Labs) were assembled onto the cleaned coverslips. A drop of immersion oil was applied to the objective lens prior to mounting the coverslip.

The instrument was calibrated before each experiment using β-amylase (56, 112, and 224 kDa) and thyroglobulin (335 and 670 kDa). For focusing, 17 μL of buffer was added to a gasket well, and the focus was automatically found using the AcquireMP software. Samples were diluted in corresponding assembly buffers to final concentrations of 100 nM. Subsequently, 3 μL of diluted protein (standard or sample) was added into the same well. After stabilization of the autofocus, 60-s movies were recorded. Each sample was measured in three independent replicates. Image processing and analysis were performed using DiscoverMP software.

### *In vitro* transposition assay

Proteins required for in vitro transposition, TnsA, TnsB, TnsC, and TnsD for TnsD-guided transposition, and TnsA, TnsB, TnsC, TniQ, and *Av*Cascade for RNA-guided transposition, were diluted to 5 μM in dilution buffer containing 25 mM Tris-HCl, pH 8.0, 500 mM NaCl, 1 mM EDTA, 1 mM DTT, and 25% glycerol. 200 nM of each protein, 100 ng pTarget, and 100 ng pDonor were added to the reaction buffer containing 26 mM HEPES (pH 7.5), 4.2 mM Tris-HCl pH 8.0, 2.1 mM DTT, 0.05 mM EDTA, 0.2 mM MgCl_2_, 28 mM NaCl, 21 mM KCl, 1.35% glycerol, 50 μg/mL BSA, and 2 mM ATP (final pH 7.5) for a total volume of 20 μL as described before. Reactions were incubated at 30 °C for 30 min, followed by supplementation with 25 mM MgOAc2 and further incubation at 30 °C for another 2 hours.

After completion, 1 μL of the reaction product was used for direct PCR analysis, 1 μL was used for quantitative PCR (qPCR), and 15 μL was used for bacterial transformation and kanamycin selection to isolate transposition products.

To test the effect of TnsD mutants, 0.78 nM variants were added to the reaction for qPCR. To test the effect of Cascade mutants, 0.62 nM variants were added to the reaction for qPCR. To test the effect of TniQ mutants, 0.39 nM variants were added to the reaction for qPCR.

### Polymerase chain reaction (PCR)

For endpoint PCR analysis, 1 μL of the completed reaction products was mixed to a final volume of 25 μL with primers for PCR reactions using Phusion Hot Start II High-Fidelity PCR Master Mix (Thermo Scientific, #F565L). PCR cycling conditions were as follows: 1 cycle, 98°C, 3 min; 30 cycles, 98°C, 10 s, 60°C, 30 s, 72°C, 5 s; 1 cycle, 72°C, 30 s.

### Quantitative PCR

Qualitative PCR (qPCR) was performed using 2 μL of the final products mixing with 900 nM forward primer, 900 nM reverse primer, and 250 nM of LE probe using TaqMan Fast Advanced Master Mix (Applied Biosystems) using Bio-Rad CFX Opus Real-Time PCR Systems. Thermal cycling conditions were as follows: 1 cycle, 50 °C, 2 min; 1 cycle, 95 °C, 1.5 min; 39 cycles, 95 °C, 10 s, 60 °C, 30 s; End.

To quantify the amount of target plasmid remaining after in vitro transposition, samples were diluted 100-fold to fall within the linear range of the standard curve. qPCR reactions contained 2 μL of diluted sample, 900 nM forward primer, 900 nM reverse primer, and 250 nM target plasmid probe, with cycling conditions of 50 °C for 2 min; 95 °C for 1.5 min; and 39 cycles of 95 °C for 10 s and 59 °C for 30 s. Insertion frequency was calculated as the copy number of inserted donor plasmid divided by the copy number of target plasmid. Sequences of all primers and probes used in this study are listed in **Supplementary Table 1**.

### Preparation of TnsC–TnsD–DNA and STC

The DNA substrate comprises seven oligonucleotides: LUEGO, *glmS*_target, *glmS*_RE_long, and polyA_RE for the RE portion, LE_target_downstream_long, target_downstream_short, and polyA_LE for the LE portion. LUEGO is a modified DNA oligo with dethiobiotin on the 5′ end, which binds to Streptavidin beads for complex assembly via a pull-down assay ^22,52^. For RE portion, LUEGO, *glmS*_target, *glmS*_RE_long, and polyA_RE were annealed at the ratio of 1:1.5:2:3 in the 10x annealing buffer (10 mM Tris pH 7.5, 50 mM NaCl, and 1 mM EDTA) to a final concentration of 12 μM. The LE portion was annealed separately to a final concentration of 12 μM using the same buffer condition.

Magnetic streptavidin beads (100 μL slurry) were washed three times with 500 μL DNA binding buffer (20 mM HEPES, pH 7.5, 250 mM KCl, 15 mM MgCl_2_, and 1 mM DTT). Subsequently, 15 μL of the 12 μM LUEGO-containing DNA substrate was diluted into 500 μL DNA binding buffer and incubated with the beads at room temperature for 30 min on an end-to-end rotator. To assemble the TnsC-TnsD-DNA complex, the beads were washed and equilibrated with protein binding buffer containing 50 mM HEPES (pH 7.5), 100 mM NaCl, 1 mM DTT, 5 mM MgCl_2_, and 0.3 mM AMP-PNP.

Next, 37.6 μL of 25 μM TnsA, 94 μL of 25 μM TnsC, and 49 μL of 19 μM TnsD were mixed with 1019.4 μL of protein binding buffer and incubated with the DNA-bound Streptavidin beads for 1 h at room temperature with end-to-end rotation. After incubation, the supernatant was removed. 37.6 μL of 20 μM TnsB, 4 μL of 12 μM LE portion, and 1 mL of protein binding buffer were added to the beads and incubated for an additional 1 h at room temperature.

Following assembly, the beads were washed three times with protein binding buffer to remove the excess proteins and DNA. The complex sample was eluted using buffer containing 50 mM biotin, 50 mM HEPES (pH 7.5), 100 mM NaCl, 1 mM DTT, 5 mM MgCl_2_, and 0.3 mM AMP-PNP. The concentration of the eluted complex was measured by Bradford assay.

Assembly of the STC followed the procedure used for the TnsC–TnsD–DNA complex, except that ATP was included in buffers and TnsA was added together with TnsB rather than with TnsC and TnsD. Two control samples were prepared. The first control sample was prepared by omitting TnsC and TnsD in the sample preparation to confirm the essential role of TnsC and TnsD in the assembly of STC. In the second control sample, TnsA was added alongside TnsC and TnsD to rule out the possibility that formation of the TnsC–TnsD–DNA complex depends on TnsA.

### Preparation of TnsB-DNA complex

The TnsB–DNA complex was assembled by incubating purified TnsB with an 80-bp RE DNA substrate containing three TnsB-binding sites, comprising 76 bp within the transposon end and 4 bp flanking DNA (**Supplementary Table 1**). Assembly was performed under diluted conditions to prevent precipitation. Briefly, 100 µL of 20 µM TnsB and 14.8 µL of 45 µM DNA was diluted into 10 mL of buffer (20 mM HEPES 7.5, 150 mM NaCl, 2 mM MgCl_2_, and 1 mM DTT) and incubated at 37 °C for 30 min. The sample was concentrated to 1 mL using a 50 kDa cutoff concentrator and subjected to Superdex 200 Increase 10/300 GL column (Cytiva). Fractions were analyzed by SDS-PAGE and agarose gels to detect TnsB and DNA, respectively. Fractions containing both components in the shifted peak were pooled and concentrated to 0.1 mg/mL.

### Preparation of *Av*Cascade–TniQ–DNA complex

The *Av*Cascade–TniQ–DNA complex was assembled on magnetic Ni–NTA beads via the N-terminal 6×His tag on Cas8. Briefly, substrate DNA was prepared by annealing oligonucleotides *Av*PSP1_90_1 and *Av*PSP1_90_2 (**Supplementary Table 1**) in 10x annealing buffer to a final concentration of 45 μM. To assemble the *Av*Cascade–DNA complex, 38 μL of 9.6 μM *Av*Cascade was incubated with 10.5 μL of 45 μM DNA in assembly buffer A containing 20 mM HEPES (pH 7.5), 150 mM NaCl, 5 mM MgCl_2_, and 1.43 mM β-mercaptoethanol at 37 °C for 30 min.

The assembled *Av*Cascade–DNA complex was then applied to 180 μL of pre-washed magnetic Ni–NTA bead slurry and incubated at room temperature for 30 min with end-to-end rotation. The beads were washed three times with assembly buffer A to remove excess unbound Cascade and DNA and subsequently equilibrated with assembly buffer B containing 20 mM Tris–HCl (pH 8.0), 100 mM NaCl, 2% (v/v) glycerol, 5 mM MgCl_2_, 1.43 mM β-mercaptoethanol, and 0.3 mM AMP-PNP.

For TniQ and TnsC binding, 28 μL of 39 μM TniQ and 185 μL of 25 μM TnsC were mixed with 912 μL of assembly buffer B and incubated with the bead-bound *Av*Cascade–DNA complex at room temperature for 1 h. Following incubation, the beads were washed three times with assembly buffer B supplemented with 20 mM imidazole and the assembled complex was eluted using assembly buffer B supplemented with 250 mM imidazole. Protein concentration was determined by Bradford assay.

### Preparation of TnsC–TniQ–DNA complex

The TnsC-TniQ-DNA complex was assembled using magnetic streptavidin beads. The DNA substrate, containing 60 bp downstream of the protospacer and a 14-bp dethiobiotinylated LUEGO sequence, was prepared by annealing three oligonucleotides: LUEGO, CQ_60_1, and LUEGO_CQ_60_2 (**Supplementary Table 1**). The DNA substrate was immobilized on magnetic streptavidin beads in DNA binding buffer (20 mM Tris-HCl, pH 7.5, 250 mM KCl, 15 mM MgCl_2_, and 1 mM DTT) and incubated at room temperature for 1 h. Excess DNA was removed by washing with CQ binding buffer (20 mM Tris-HCl, pH 8.0, 100 mM NaCl, 1% glycerol, 5 mM MgCl_2_, and 0.3 mM AMP-PNP). For complex assembly, 20 μL of 39 μM TniQ and 97.2 μL of 25 μM TnsC were diluted into 1054.8 μL of CQ binding buffer and incubated with the DNA-bound beads for 1 h at room temperature. The beads were washed three times with CQ binding buffer to remove unbound proteins, and the assembled complex was eluted using CQ binding buffer supplemented with 50 mM biotin.

### Electron microscopy

Aliquots of 3 μL of the TnsC–TnsD–DNA complex (∼0.5 mg/mL) were applied to glow-discharged Quantifoil holey carbon grids (R1.2/1.3, 300 mesh). Samples were applied three times^53^, with blotting for 2.5 s at blot force 2 after each application, before plunge-freezing into liquid ethane using a Thermo Fisher Scientific Vitrobot Mark IV.

Cryo-EM data were collected with a Titan Krios G4 microscope (FEI) operated at 300 kV. Images were acquired using EPU at a nominal magnification of 105,000×, resulting in a calibrated physical pixel size of 0.822 Å/pixel, with a defocus range of 0.8–2.0 μm. Movies were recorded on a K3 direct electron detector in super-resolution mode at the end of a GIF Quantum energy filter operated with a 20 eV slit width. A dose rate of 20 electrons per pixel per second and an exposure time of 2.01 s were used, resulting in 40 frames per movie and a total accumulated dose of ∼59.2 e⁻/Å^2^. After grid screening, two grids were selected for data collection, yielding a total of 5,486 micrographs for structure determination of the TnsC_3_–TnsD_1_–DNA_1_ complex. To address preferred orientation in the TnsC_7_–TnsD_1_–DNA_1_ dataset, an additional 6,164 micrographs were collected using a 30° stage tilt.

For the STC complex, 0.5 mg/mL samples were applied to glow-discharged Quantifoil holey carbon and UltrAu grids. After grid screening, one Quantifoil and one UltrAuFoil holy carbon grid were selected for data collection under similar conditions used for the TnsC_3_–TnsD_1_–DNA_1_ complex, yielding 2,883 and 13,097 micrographs, respectively.

The TnsB–DNA complex (0.1 mg/mL) was applied to glow-discharged Quantifoil grids (R2/2, 300 mesh) coated with a 2-nm carbon layer. Data were collected under conditions similar to those used for the TnsC_3_–TnsD_1_–DNA_1_ complex, yielding 1,119 micrographs.

For the *Av*Cascade–TniQ–DNA and TniQ–TnsC–DNA complexes (∼0.5 mg/mL), cryo-EM grid preparation and data collection conditions were similar to those used for the TnsC_3_–TnsD_1_–DNA_1_ sample, except that the Cascade–TniQ–DNA sample was applied only once prior to blotting. In total, 8,247 micrographs were collected for the Cascade–TniQ–DNA dataset and 2,458 micrographs for the TniQ–TnsC–DNA dataset.

### Image processing

For the TnsC–TnsD–DNA dataset, movie frames were imported to cryoSPARC ^54^ live for on-the-fly processing during data collection. Motion correction was performed using patch motion correction with a binning factor of 2, and Contrast transfer function (CTF) parameters were estimated using Patch CTF. Particles were auto-picked without a template to generate 2D averages. The auto-picked and extracted particles were screened by 2D classification to exclude false and bad particles that fall into 2D averages with poor features. A subset of the particles (200,000 particles) was used to generate initial models in cryoSPARC. The particles were further classified by heterogeneous refinement and 2D classification. Particles with clear features for TnsC_3_–TnsD_1_–DNA_1_ complex and TnsC_7_–TnsD_1_–DNA_1_ after 2D classification were used for Topaz training separately^55^. For TnsC_3_–TnsD_1_–DNA_1_ complex, 788,816 particles were picked using the Topaz model. After 2D classification, 713,441 particles were selected for further analysis. Two major classes were identified from 3D classification, one TnsC_3_–TnsD_1_–DNA_1_ complex (173,082 particles), and the other TnsC_7_–TnsD_1_–DNA_1_ complex (294,575 particles). Similar procedures were done for TnsC_7_–TnsD_1_–DNA_1_. However, the TnsC_7_–TnsD_1_–DNA_1_ complex showed stretched features indicating preferred orientation. Further 3D classification of the particles of TnsC_3_–TnsD_1_–DNA_1_ complex resulted in a TnsC_4_–TnsD_1_–DNA_1_ complex with a final data set of 11,240 particles and a TnsC_3_–TnsD_1_–DNA_1_ complex with a final data set of 91,633 particles. After non-uniform refinement, we obtained a TnsC_3_–TnsD_1_–DNA_1_ structure at 3.08 Å resolution and a TnsC_4_–TnsD_1_–DNA_1_ at 3.82 Å resolution. The resolution was estimated based on the fourier shell correlation (FSC) = 0.143 criterion. The map was postprocessed using DeepEMhancer ^56^ to reduce noise levels. This postprocessed map was used for model building and refinement. A similar workflow was used for TnsC_7_–TnsD_1_–DNA_1_ complex with stage tilt and TniQ–TnsC–DNA complex.

For the STC dataset, two rounds of Topaz training were performed to generate a Topaz model. 1,851,842 particles were initially selected. After three rounds of heterogeneous refinement and one round of 3D classification, 51,307 particles were retained, yielding a 3.19 Å density map following non-uniform refinement. A mask was applied to the asymmetric region of transposon end for local refinement, improving local resolution for model building.

The well-trained Topaz model for STC was also applied to pick particles from the TnsC–TnsD–DNA datasets described above; however, no particles were retained after 2D classification. Topaz models trained on the TnsC_3_–TnsD_1_–DNA_1_ and TnsC_7_–TnsD_1_–DNA_1_ complexes likewise failed to identify STC particles in the STC datasets.

For the TnsB–DNA dataset, particles were initially selected by blob and template picking to generate training data for Topaz. A total of 295,815 particles were selected. After 2D classification and heterogeneous refinement, 187,897 particles were retained for 3D classification. Of six classes, two (54,538 particles) represented a TnsB–DNA complex containing four TnsB subunits. Particles were re-extracted from micrographs with a bin-1 setting, yielding 53,796 particles for final non-uniform refinement and a 3.9 Å density map.

For the Cascade–TniQ–DNA dataset, similar workflow was used to generate the Topaz model. 1,404,457 particles were picked using the trained picking model, and 1,072,190 particles were further used after 2D classification. Three classes were identified after heterogeneous refinement. Class 0 (491,063 particles) showed density around the PAM-distal area, Class 1 (421,371 particles) had no extra density around the PAM-distal area, and Class 2 (159,756 particles) with bad or false particles. Particles from class 1 were further classified using heterogenous refinement. The resulting 396,010 particles were used to reconstruct a map for *Av*Cascade–DNA-1 with flexible Cas6 at 2.64 Å. Particles in class 0 were further classified using heterogenous refinement for 2 rounds, resulting in a final particle set of 326,574 particles. A mask was generated around the PAM-distal area for focused 3D classification, which resulted in two groups of particles, one group for *Av*Cascade–TniQ–DNA (42,834 particles) and one for *Av*Cascade–DNA-2 with flexible Cas8 (49,799 particles). Using non-uniform Refinement, we obtained a 3.16 Å density map for *Av*Cascade–TniQ–DNA and 3.01 Å density map for *Av*Cascade–DNA-2.

Data processing for the TniQ–TnsC–DNA complex followed a workflow similar to that used for the TnsC–TnsD–DNA dataset. A total of 332,402 particles were picked using Topaz, of which 288,180 were retained after 2D classification. Three rounds of heterogeneous refinement revealed two particle populations: a TnsC–DNA complex (64,288 particles) and a TniQ–TnsC–DNA complex (179,796 particles). Particles corresponding to the TnsC–DNA complex were subjected to non-uniform refinement, yielding a 3.78 Å density map. For the TniQ–TnsC–DNA complex, subsequent 3D classification further refined the dataset to 22,162 particles, resulting in a 3.56 Å density map. Additional particle populations, including species with two TnsC rings on a single DNA, as well as TnsC dimers and heptamers, were observed. Reconstructions of the TnsC dimer and trimer yielded density maps at 6.70 Å and 6.60 Å resolution, respectively.

### Model building and refinement

Initial structures of TnsC_7_–TnsD_1_–DNA_1_, TniQ, *Av*Cascade–DNA, and STC were predicted by Alphafold^57^. The models were first fitted into the maps as a rigid-body in UCSF ChimeraX^58^ and then manually adjusted in COOT^59^. The resolution in TnsC_7_–TnsD_1_–DNA_1_, STC, and *Av*Cascade–DNA–TniQ allowed us to distinguish between purines and pyrimidines, which allowed us to confidently assign the sequences in the models. The models were then fitted in COOT with all molecule self-restrains to maintain hydrogen bonds and stacking interactions between bases. The DNA sequence in TnsC_3_–TnsD_1_–DNA_1_ was assigned based on the DNA sequence and density features in the two DNA segments (**Supplementary Fig. 3**). Finally, refinement of the structure models against corresponding maps were performed using phenix.real_space_refine tool in Phenix^60^.

## Supporting information

Extended Data Tabel 1

## Acknowledgements

We thank Frank Vago and Thomas Klose for assistance in cryo-EM data collection and Steven Wilson for computation. We are grateful to Dr. Andrew D. Mesecar and Uttara Jayashankar at Purdue University for help with mass photometry. We thank Purdue Proteomics facility and its staff at the Bindley Bioscience center for the proteomics support. This work was supported by the NIH grant R35GM158248, NSF CAREER 2339799, and a Core Pilot grant from the Indiana Clinical and Translational Sciences Institute to L.C.

## Author contributions

S.W. assembled all complexes for cryo-EM analysis, prepared cryo-EM grids, collected and processed cryo-EM data, built and analyzed structural models, prepared figures, and drafted the manuscript. S.W. cloned pTarget, designed DNA substrates and structure-guided mutants for TnsD, *Av*Cascade, and TniQ, generated and purified *Av*Cascade and TniQ mutant proteins, performed qPCR-based activity assays and BamHI cleavage assays, and prepared samples for mass spectrometry and mass photometry. R.S. constructed plasmids for TniQ, *Av*Cascade, pDonor, TnsD mutants, and pTarget mutants; purified *Av*CAST components, including TnsA, TnsB, TnsC, TnsD, TniQ, *Av*Cascade, and TnsD mutant proteins; prepared proteins used for complex formation; performed in vitro transposition assays by direct PCR for TnsD and pTarget mutants, and tested the minimal LE and RE requirements for transposition. L.C. supervised the study, secured funding, and edited and revised the manuscript with input from all authors.

## Competing interests

The authors declare no competing interests.

## Data, code, and materials availability

Cryo-EM reconstructions and atomic coordinates generated in this study have been deposited in the Electron Microscopy Data Bank and Protein Data Bank, respectively. The accession numbers and associated structural data will be made publicly available upon publication of the peer-reviewed article.

---

**Extended Data Fig. 1.**
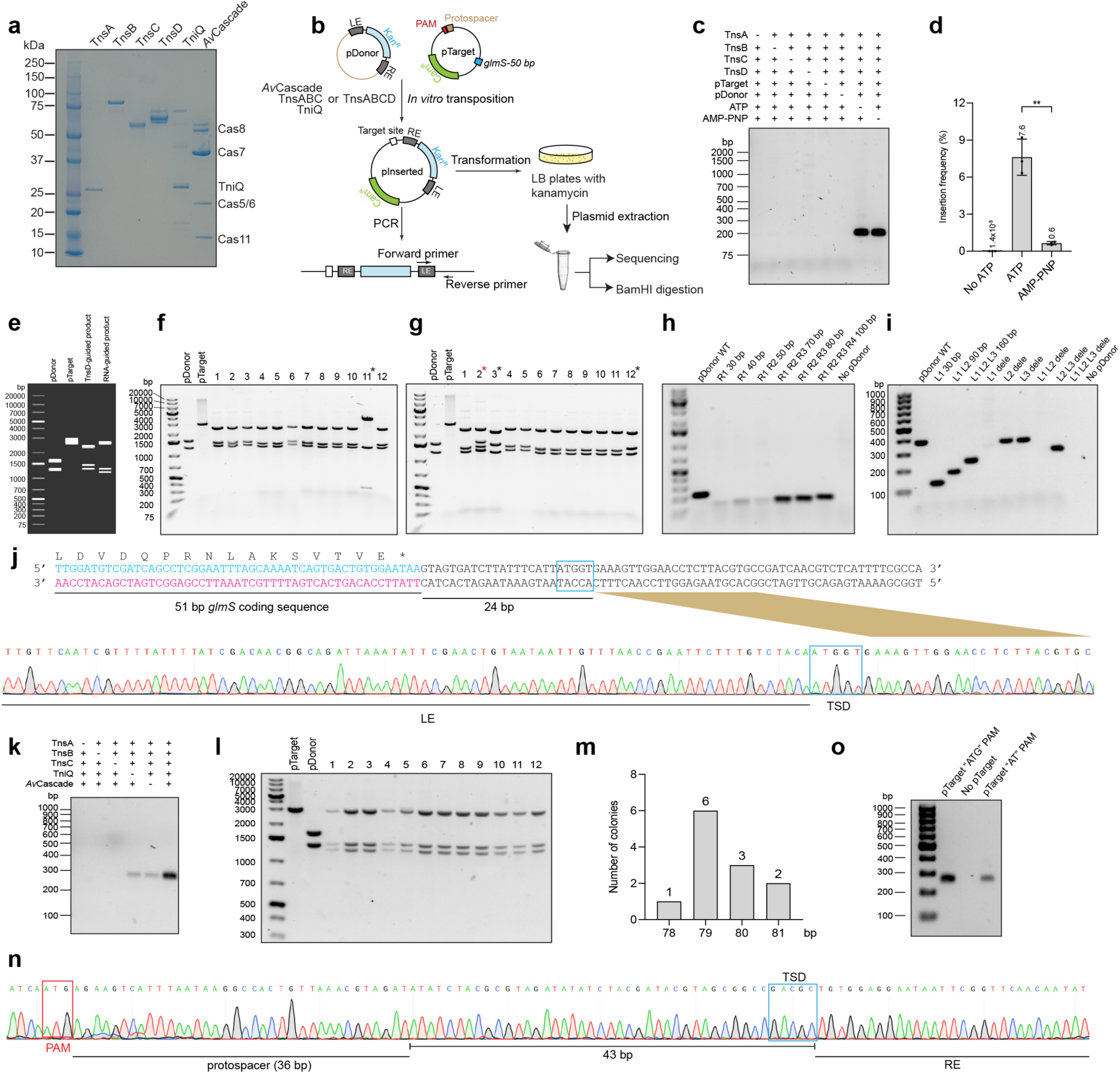
Biochemical reconstitution of TnsD-guided and RNA-guided transposition by *Av*CAST. **a**, SDS–PAGE analysis of purified *Av*CAST components used for biochemical reconstitution. **b**, Schematic of the *in vitro* transposition assay, including donor and target plasmids, transformation-based recovery of product plasmids, PCR detection, sequencing and BamHI digestion analysis. **c**, Direct PCR detection of transposition products with the indicated components and nucleotide conditions. **d**, Quantification of insertion frequency under no-ATP, ATP and AMP-PNP conditions. **e**, BamHI digestion of a simulated transposition product, used as a reference for interpreting recovered products. **f,g**, BamHI digestion analysis of plasmids recovered from TnsD-guided transposition products generated in the presence of ATP (**f**) or AMP-PNP (**g**). Black asterisks indicate off-target simple insertions; the red asterisk indicates an on-target cointegrate product. **h,i**, Truncation analysis of RE (**h**) and LE (**i**) substrates, defining the minimal TnsB-binding-site requirements for transposition. **j**, Sanger sequencing of a representative TnsD-guided insertion product, showing insertion 24 bp downstream of the *glmS* stop codon and RE positioned proximal to the target site. **k**, Direct PCR detection of RNA-guided transposition products. **l**, BamHI digestion analysis of plasmids recovered from RNA-guided transposition products. **m**, Distribution of RNA-guided insertion positions downstream of the PAM. **n**, Sanger sequencing of a representative RNA-guided insertion product. **o**, PCR analysis comparing target plasmids containing an ATG PAM, no pTarget, or an AT PAM.

**Extended Data Fig. 2.**
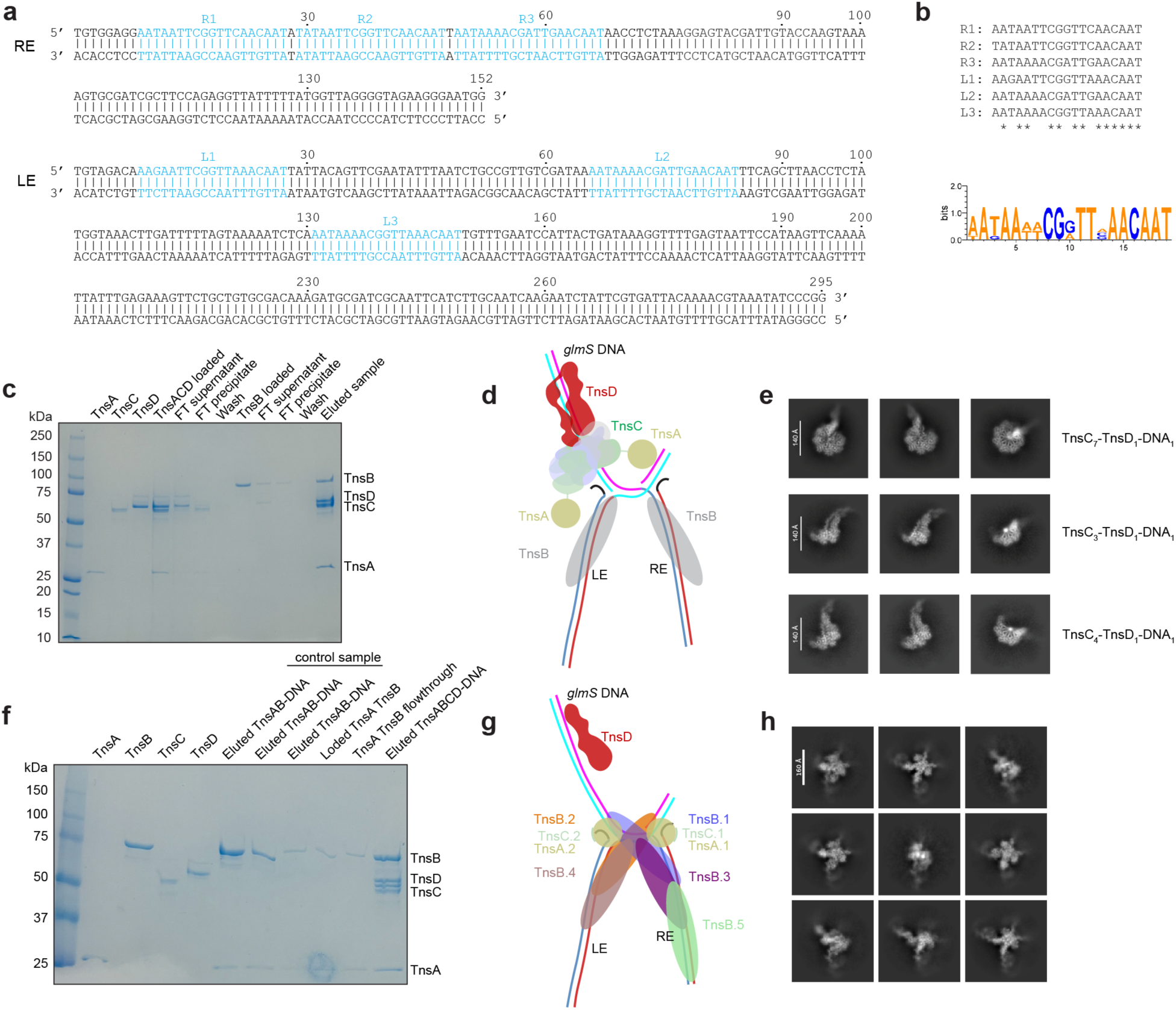
Transposon-end organization and assembly of *Av*CAST TnsC–TnsD–DNA and STC complexes. **a**, Sequences of the *Av*CAST RE and LE, with TnsB-binding sites indicated. **b**, Alignment and sequence logo of TnsB-binding sites from RE and LE. **c**, SDS–PAGE analysis of affinity-purified complexes assembled with TnsA, TnsB, TnsC, TnsD and DNA in the presence of AMP-PNP. **d**, Schematic of the AMP-PNP assembly outcome. Although TnsA, TnsB, TnsC and TnsD were present in the eluted fraction, cryo-EM resolved only TnsC–TnsD–DNA assemblies, indicating that TnsA and TnsB were not stably incorporated into the resolved particles. **e**, Representative 2D class averages of TnsC_7_–TnsD_1_–DNA_1_, TnsC_3_–TnsD_1_–DNA_1_ and TnsC_4_–TnsD_1_–DNA_1_ particles obtained from the AMP-PNP assembly. **f**, SDS–PAGE analysis of complexes assembled under ATP-containing conditions. **g**, Schematic of the ATP-containing assembly outcome, in which an asymmetric STC containing TnsA, TnsB and the C-terminal region of TnsC was resolved; TnsD was detected biochemically but not resolved by cryo-EM. **h**, Representative 2D class averages of the STC particles.

**Extended Data Fig. 3.**
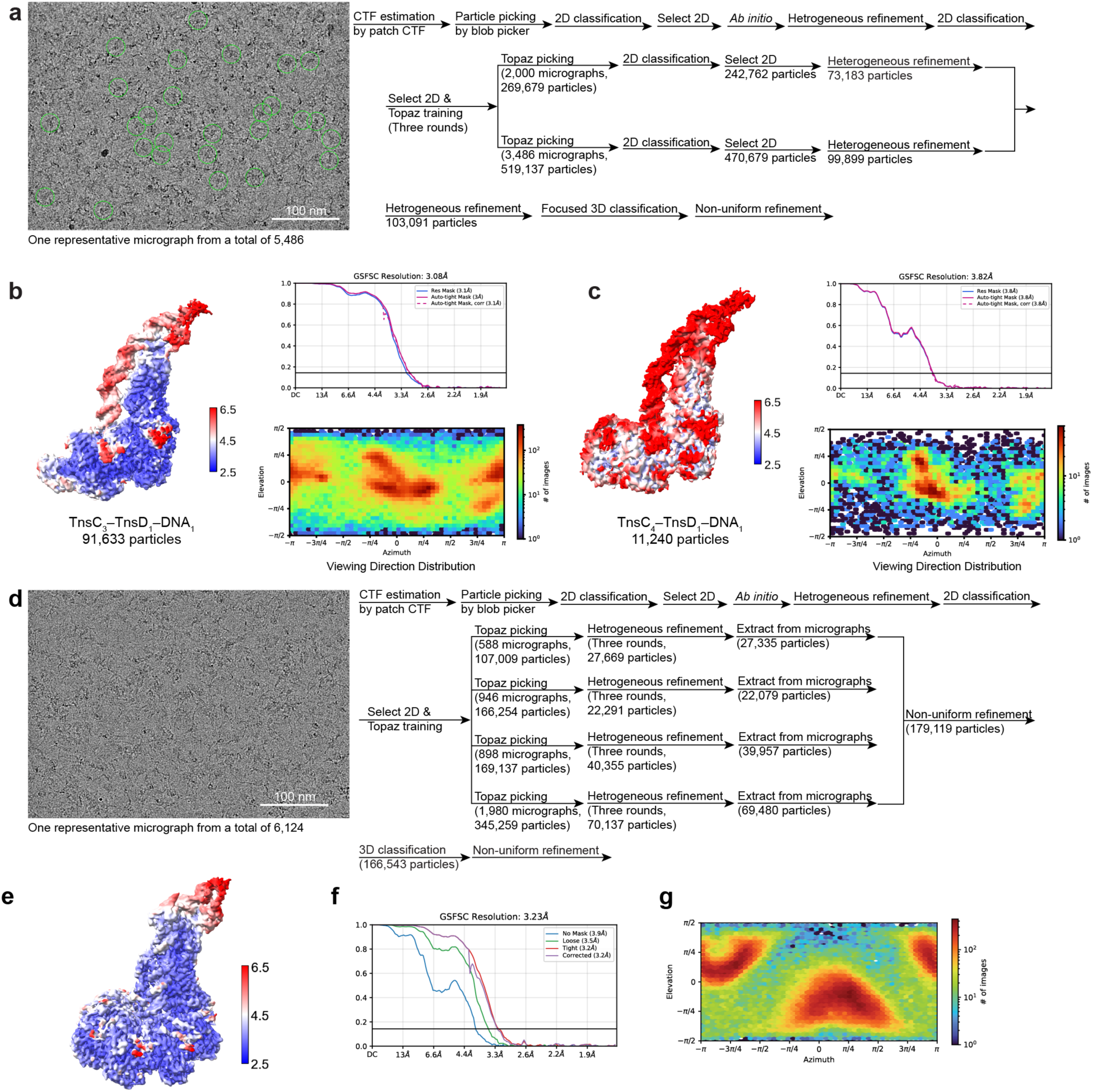
Cryo-EM processing of TnsD–TnsC assemblies on *glmS* DNA. **a**, Representative cryo-EM micrograph and data-processing workflow for the TnsC_3_–TnsD_1_–DNA_1_ and TnsC_4_–TnsD_1_–DNA_1_ assemblies. **b**, Local-resolution map, gold-standard FSC curve and viewing-direction distribution for the TnsC_3_–TnsD_1_–DNA_1_ reconstruction. **c**, Local-resolution map, gold-standard FSC curve and viewing-direction distribution for the TnsC_4_–TnsD_1_–DNA_1_ reconstruction. **d**, Representative cryo-EM micrograph and processing workflow for the tilted-stage TnsC_7_–TnsD_1_–DNA_1_ dataset, collected to reduce preferred orientation. **e**, Local-resolution map of the TnsC_7_–TnsD_1_–DNA_1_ reconstruction. **f,g**, Gold-standard FSC curve (**f**) and viewing-direction distribution (**g**) for the TnsC_7_–TnsD_1_–DNA_1_ reconstruction.

**Extended Data Fig. 4.**
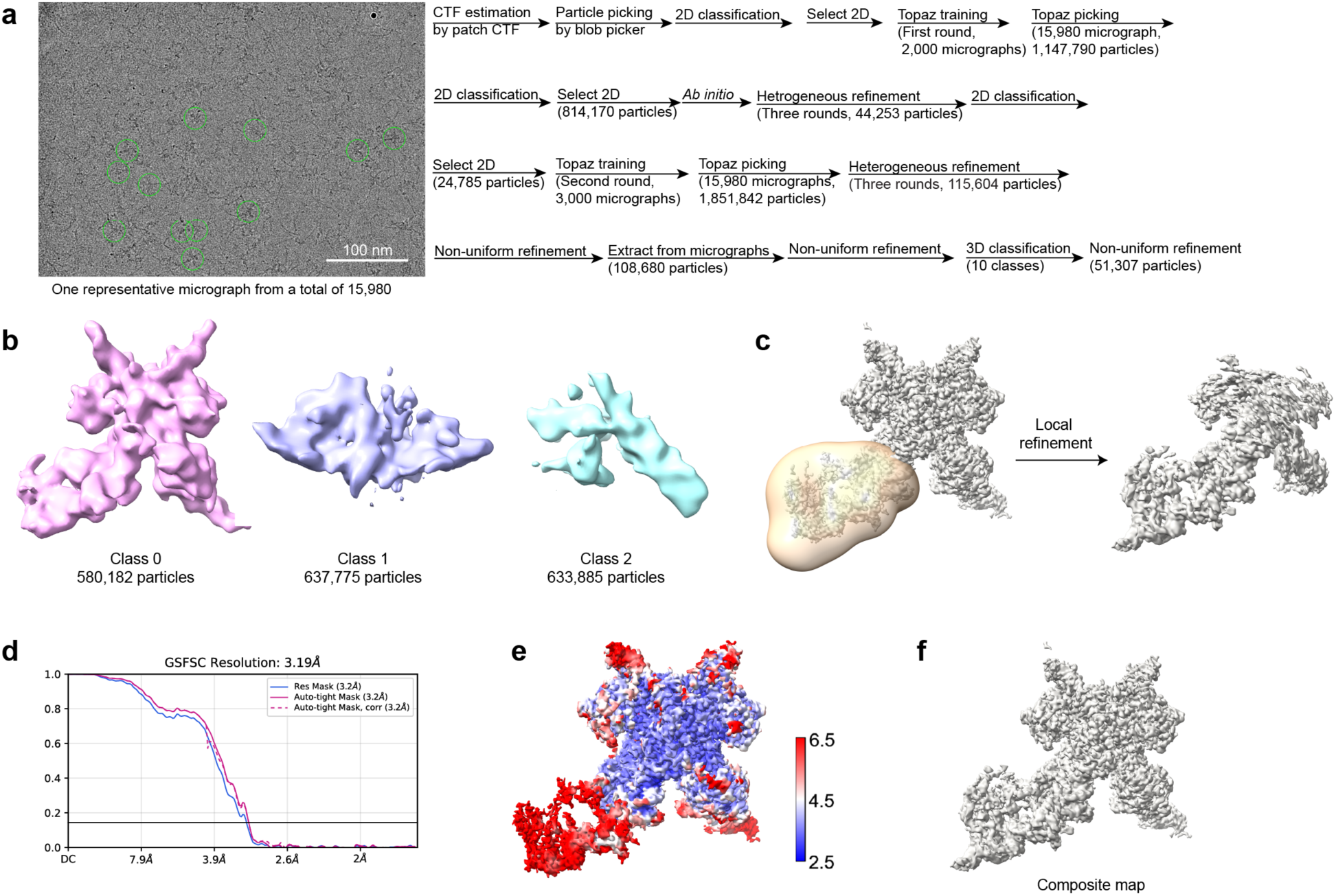
Cryo-EM processing of the *Av*CAST STC. **a**, Representative cryo-EM micrograph and data-processing workflow for the *Av*CAST STC assembled under ATP-containing conditions. **b**, Three-dimensional classification of STC particles, showing representative classes. **c**, Local refinement strategy used to improve the STC reconstruction. **d**, Gold-standard FSC curve for the final STC reconstruction. **e**, Local-resolution map of the final STC reconstruction. **f**, Composite cryo-EM map of the STC generated by combining the pre- and post-local-refinement maps shown in **c.**

**Extended Data Fig. 5.**
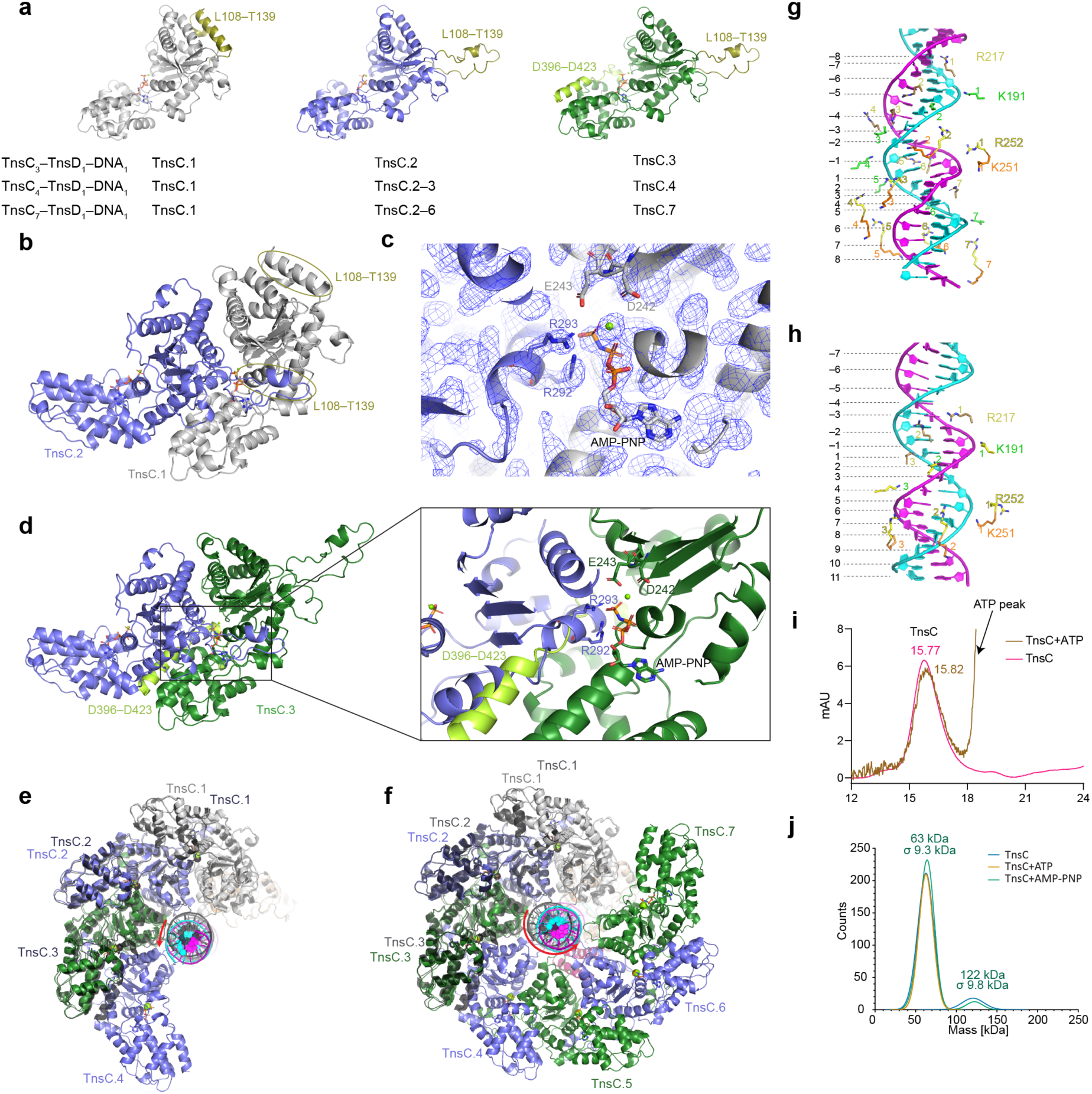
Structural basis for stepwise TnsC oligomerization on target DNA. a, Conformations of individual TnsC protomers in the TnsC–TnsD–DNA assemblies. The L108–T139 region and C-terminal self-inhibitory segment are indicated. **b**, Interface between TnsC.1 and TnsC.2 in the TnsC_3_–TnsD_1_–DNA_1_ assembly, highlighting the conformational difference of the L108–T139 region. **c**, Close-up view of the AMP-PNP-binding site in TnsC.1 in the TnsC_3_–TnsD_1_–DNA_1_ assembly, showing engagement by the arginine fingers from TnsC.2. **d**, Model of an additional TnsC protomer docked onto TnsC.3 in the TnsC_3_–TnsD_1_–DNA_1_ assembly, showing that the C-terminal self-inhibitory segment of TnsC.3 sterically clashes with the incoming protomer and blocks further TnsC recruitment. **e**, Comparison of TnsC–DNA contacts in the TnsC_3_–TnsD_1_–DNA_1_ and TnsC_4_–TnsD_1_–DNA_1_ structures, showing movement of DNA towards TnsC.4. **f**, Comparison of TnsC–DNA contacts in the TnsC_3_–TnsD_1_–DNA_1_ and TnsC_7_–TnsD_1_–DNA_1_ structures, showing movement of DNA towards TnsC.4–7. **g,h**, DNA interactions with positively charged TnsC residues in the TnsC_3_–TnsD_1_–DNA_1_ (**g**) and TnsC_7_–TnsD_1_–DNA_1_ (**h**) assemblies, showing a shift from major-groove-associated contacts in the early complex to minor-groove-associated contacts in the heptameric complex.. **i**, Size-exclusion chromatography profiles of purified TnsC in the absence or presence of ATP or AMP-PNP. **j**, Mass photometry analysis of purified TnsC under the indicated nucleotide conditions.

**Extended Data Fig. 6.**
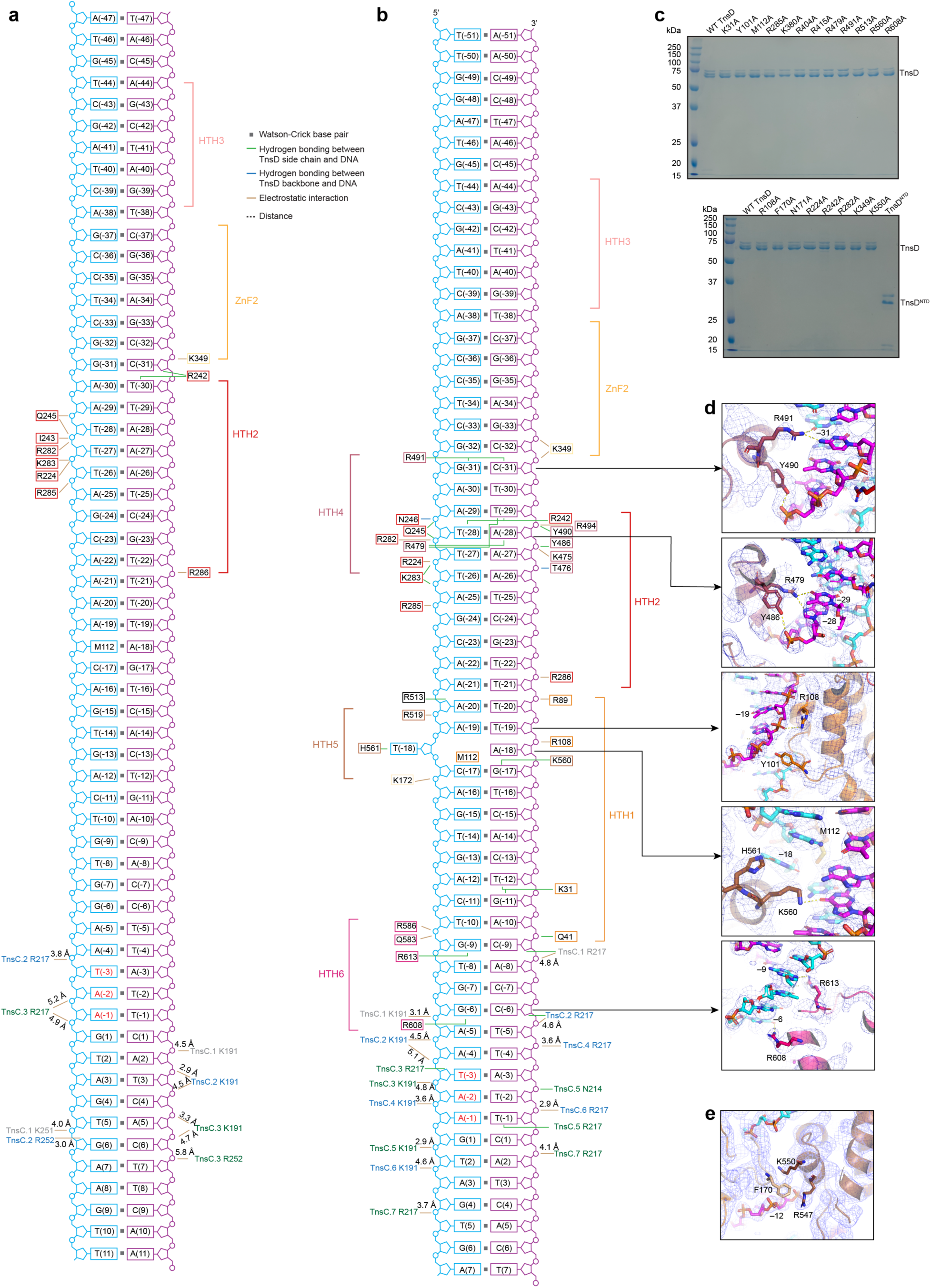
Structural and biochemical analysis of TnsD–*glmS* DNA recognition. **a,b**, Schematic of TnsD and TnsC contacts with *glmS* target DNA in the TnsC_3_–TnsD_1_–DNA_1_ (**a**) and TnsC_7_–TnsD_1_–DNA_1_ (**b**) assemblies. Hydrogen bonds, electrostatic interactions and measured distances are indicated. **c**, SDS–PAGE analysis of purified TnsD variants used for transposition assays. **d**, Representative density and model views of base-specific TnsD–DNA interactions in the TnsC_7_–TnsD_1_–DNA_1_ structure. **e**, Representative density and model view of an intramolecular NTD–CTD interface in TnsD, with F170, K550 and R547 contributing to interdomain packing.

**Extended Data Fig. 7.**
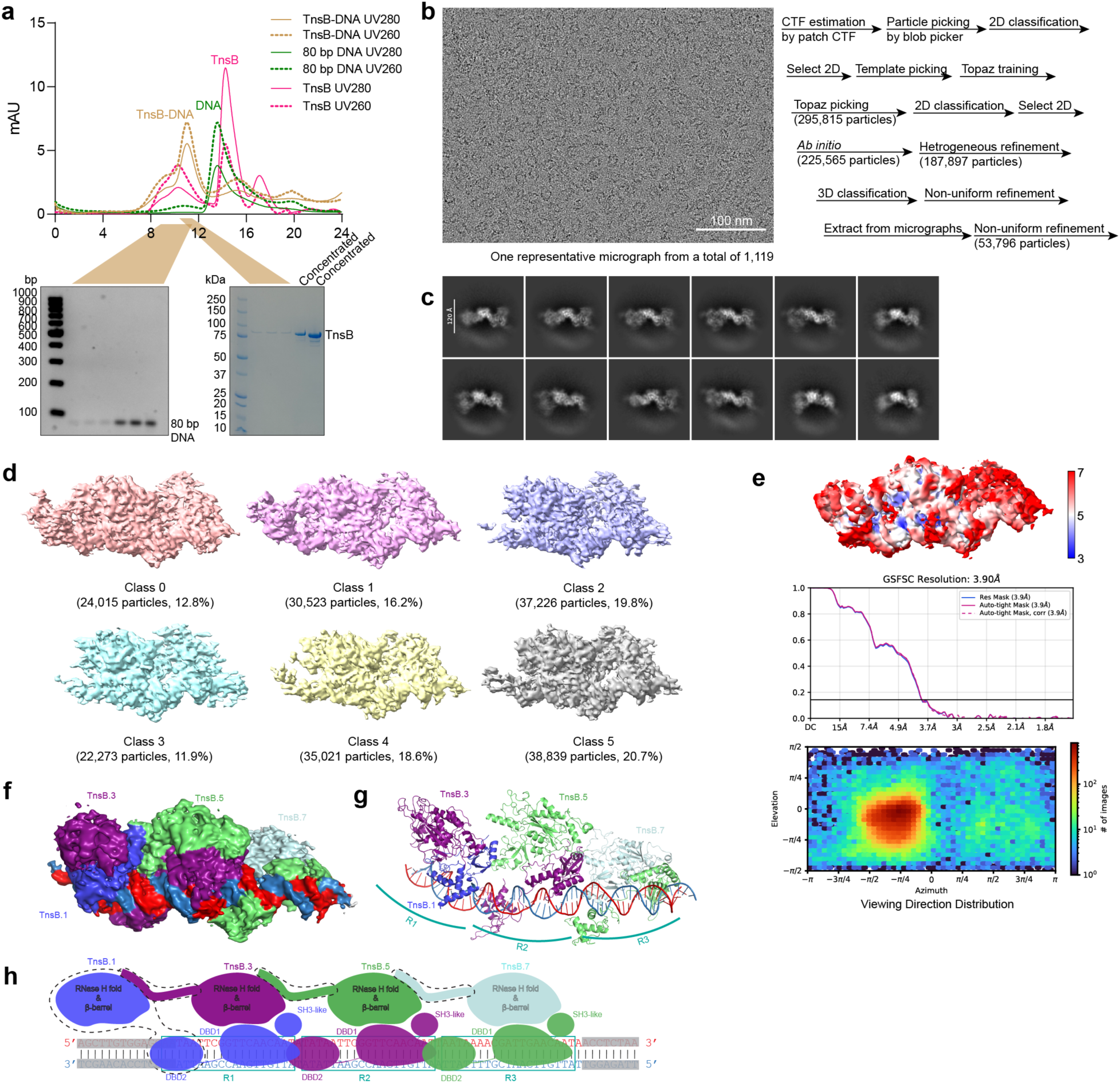
Cryo-EM structure of the TnsB–RE DNA complex. **a**, Size-exclusion chromatography profile of the TnsB–RE DNA complex. Corresponding DNA and protein sample analysis are shown below. **b**, Representative cryo-EM micrograph and data-processing workflow for the TnsB–RE DNA complex. **c**, Representative 2D class averages. **d**, Three-dimensional classification of TnsB–RE DNA particles. **e**, Local-resolution map, gold-standard FSC curve and viewing-direction distribution for the final TnsB–RE DNA reconstruction. **f,g**, Cryo-EM density map (**f**) and atomic model (**g**) of the TnsB–RE DNA complex. **h**, Schematic of TnsB protomer organization on RE DNA, showing TnsB-binding sites R1–R3 and the arrangement of TnsB domains.

**Extended Data Fig. 8.**
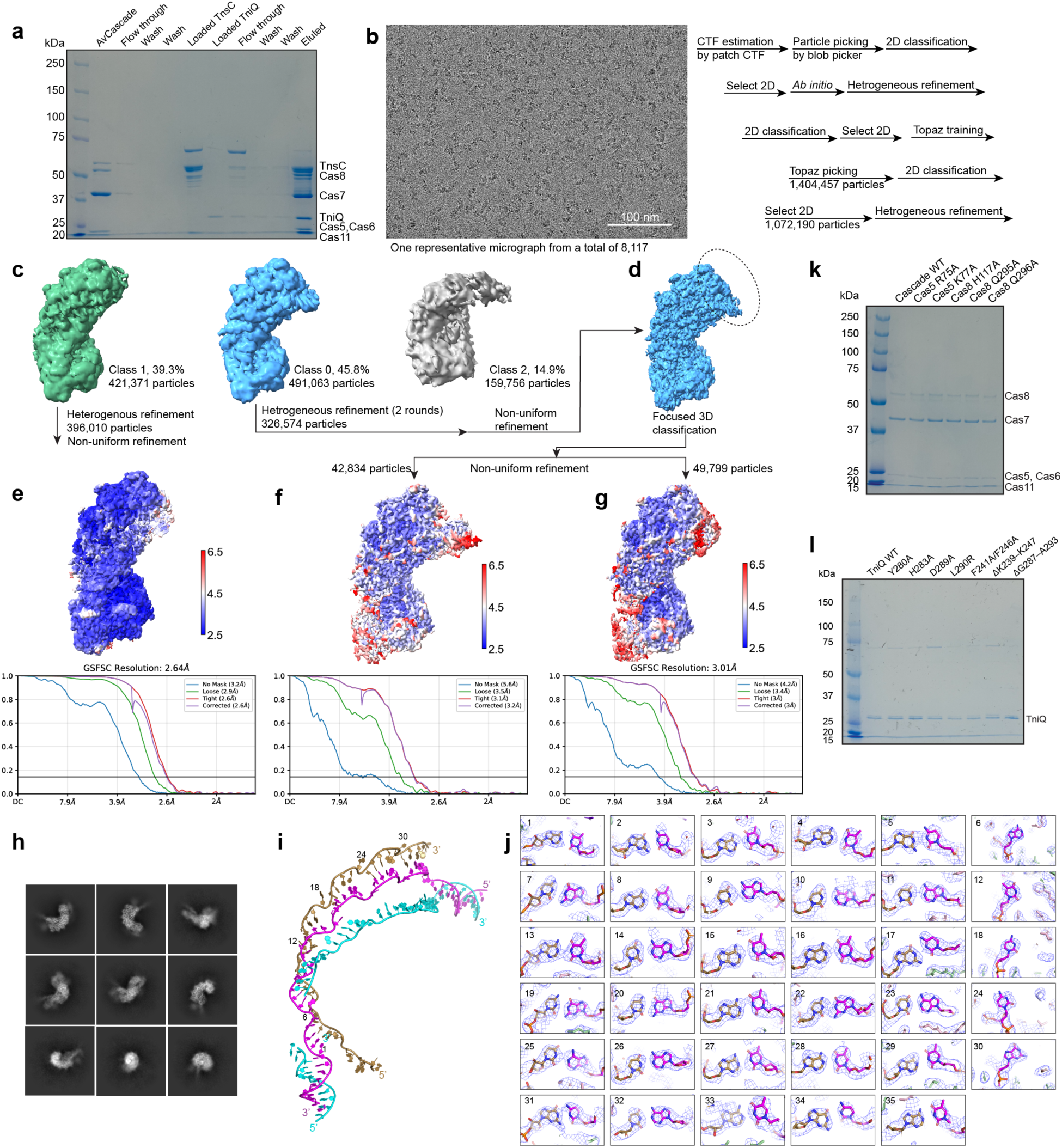
Cryo-EM processing of *Av*Cascade–DNA and *Av*Cascade–TniQ–DNA complexes. **a**, SDS–PAGE analysis of affinity-purified *Av*Cascade–DNA and *Av*Cascade–TniQ–DNA assemblies. **b**, Representative cryo-EM micrograph and data-processing workflow for *Av*Cascade–DNA and *Av*Cascade–TniQ–DNA particles. **c**, Three-dimensional classification of *Av*Cascade-containing particles, showing classes used for subsequent refinement. **d**, Focused classification of the TniQ-containing region. **e–g**, Local-resolution maps and gold-standard FSC curves for the refined *Av*Cascade–DNA and *Av*Cascade–TniQ–DNA reconstructions. **h**, Representative 2D class averages of *Av*Cascade–TniQ–DNA particles. **i**, Model of the crRNA–target DNA heteroduplex and displaced non-target strand in the *Av*Cascade–TniQ–DNA structure. **j**, Representative density for the crRNA–target DNA duplex, supporting sequence assignment. **k**, SDS–PAGE analysis of purified *Av*Cascade variants used for transposition assays. **l**, SDS–PAGE analysis of purified TniQ variants used for transposition assays.

**Extended Data Fig. 9.**
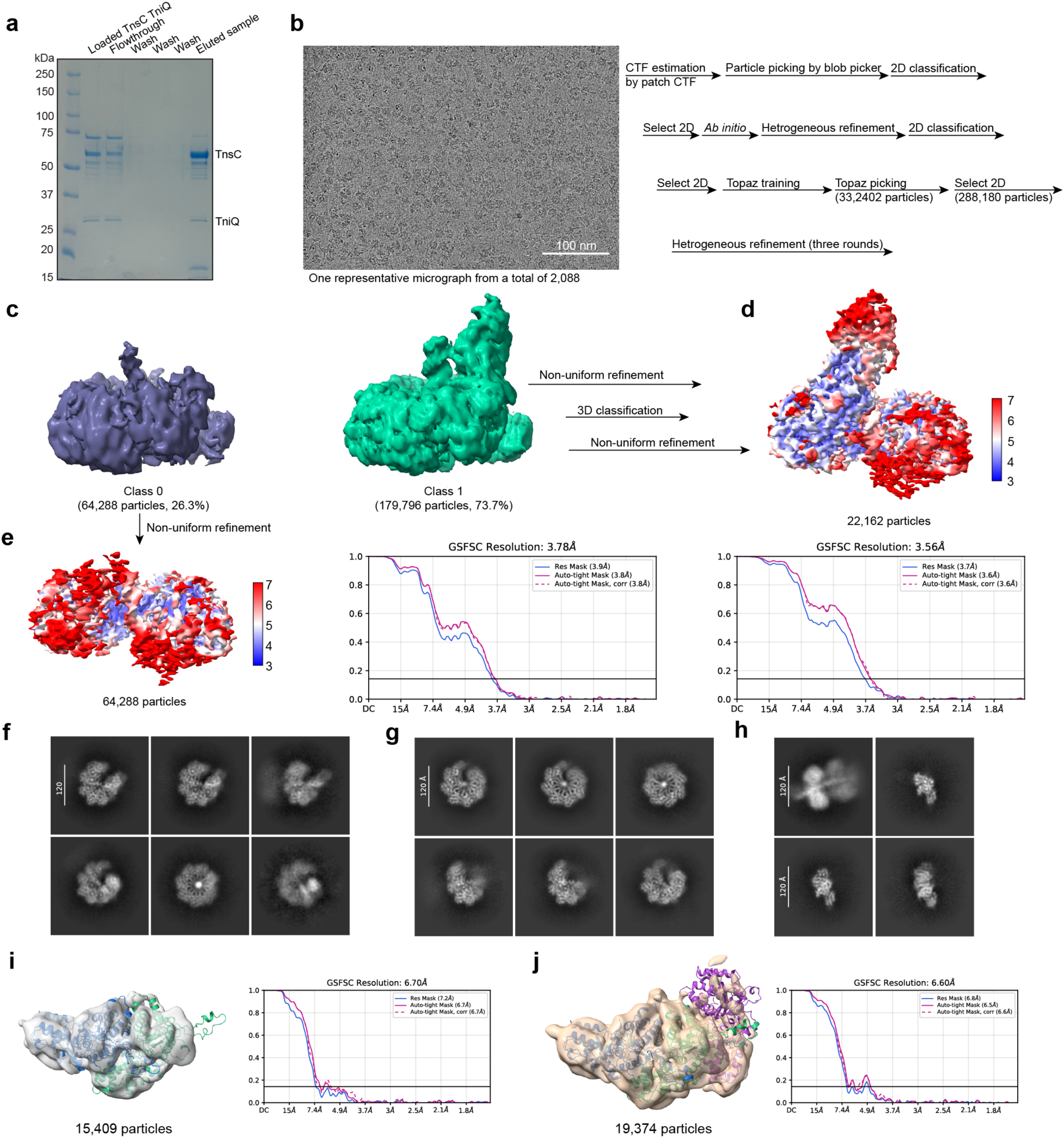
Cryo-EM processing of the TniQ–TnsC–DNA assembly. **a**, SDS–PAGE analysis of affinity-purified TniQ–TnsC–DNA assembly. **b**, Representative cryo-EM micrograph and data-processing workflow for the TniQ–TnsC–DNA assembly. **c**, Three-dimensional classification of TniQ–TnsC–DNA particles. **d,e**, Local-resolution maps and gold-standard FSC curves for the refined TniQ–TnsC–DNA **(d)** and TnsC–DNA **(e)** reconstructions, respectively. **f–h**, Representative 2D class averages of TnsC-containing particles, including TniQ-bound particles **(f)**, particles lacking visible TniQ density **(g)**, and additional minor classes **(h)**. **i,j**, Low-resolution reconstructions of TnsC-containing dimeric **(i)** and trimeric **(j)** classes observed during 3D classification.

**Extended Data Fig. 10.**
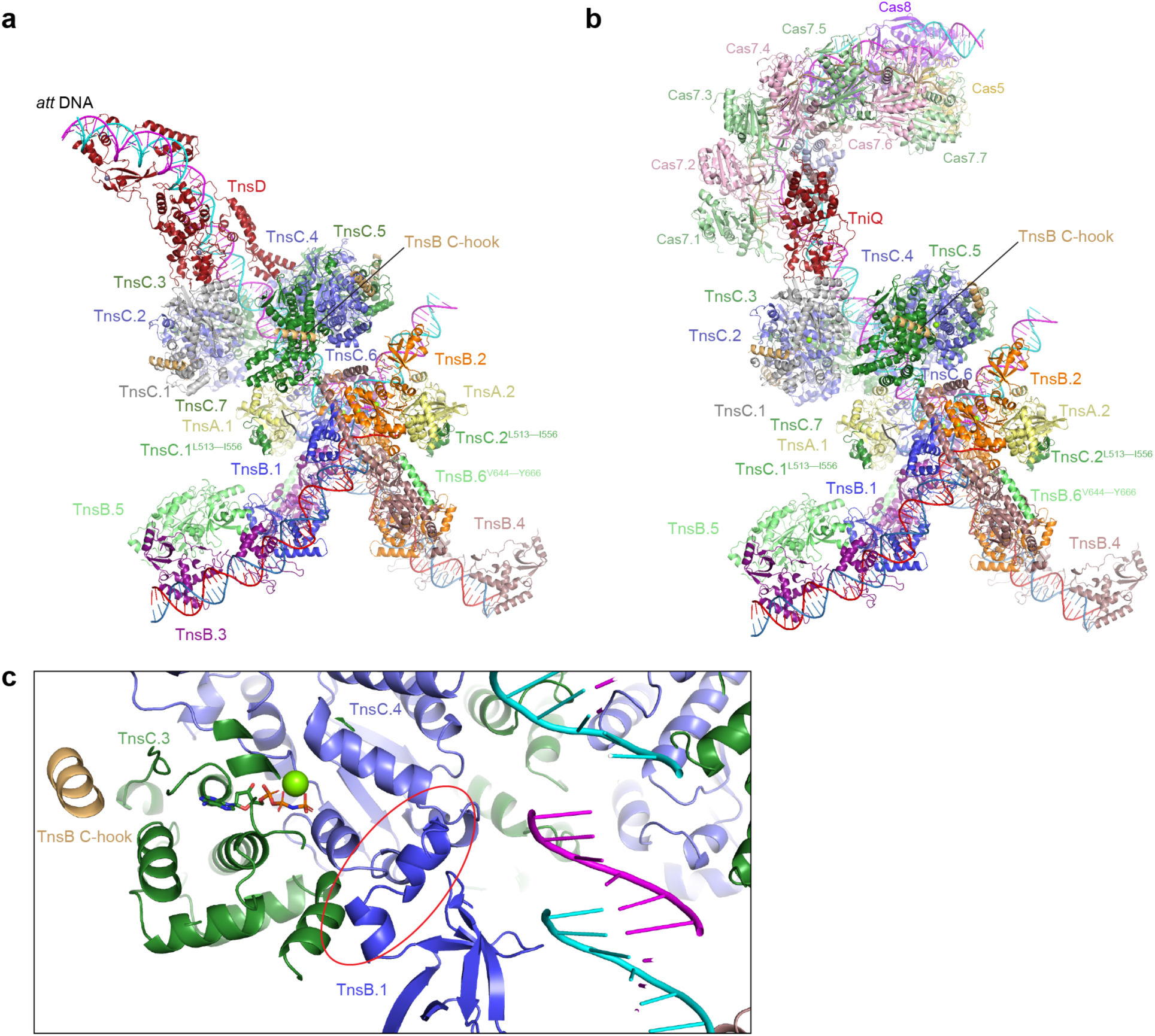
Composite models linking *Av*CAST targeting assemblies to the STC. **a**, Composite model of the TnsD-guided targeting assembly and STC, generated by superimposing target DNA from the TnsC_7_–TnsD_1_–DNA_1_ and STC structures. **b**, Composite model of the RNA-guided targeting assembly and STC, generated by superimposing shared TniQ–DNA and target-DNA elements from the *Av*Cascade–TniQ–DNA, TniQ–TnsC–DNA and STC structures. **c**, Close-up view of the composite STC model, showing that TnsB.1 would clash with the TnsC ring via TnsC.4. The TnsB C-hook was modeled based on an AlphaFold prediction of the TnsB–TnsC complex.

## Notes

### Competing Interest Statement

The authors have declared no competing interest.

