## Supplementary Information for "DNA remodeling couples target recognition to directional transposition in a Tn7-like CAST"

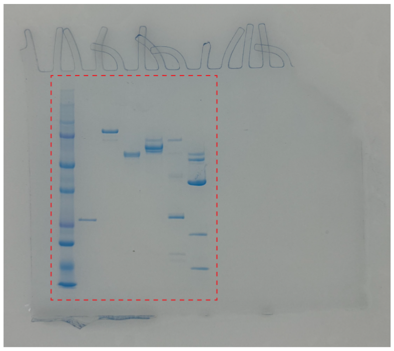
Extended Data Fig. 1a

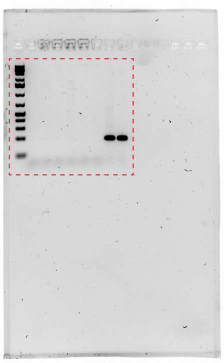
Extended Data Fig. 1c

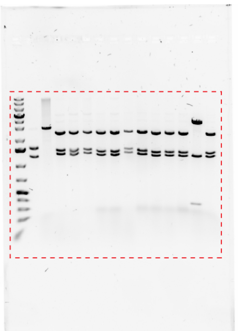
Extended Data Fig. 1f

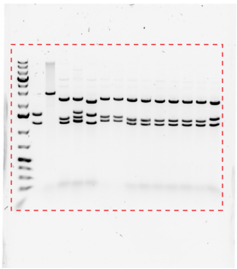
Extended Data Fig. 1g

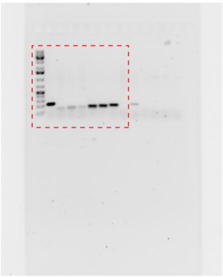
Extended Data Fig. 1h

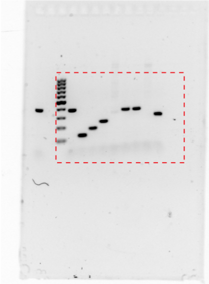
Extended Data Fig. 1i

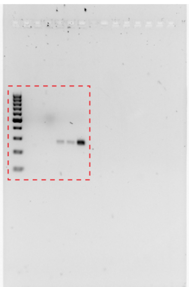
Extended Data Fig. 1k

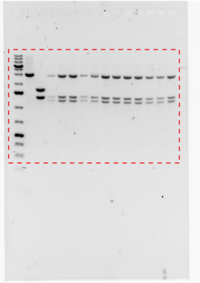
Extended Data Fig. 1l

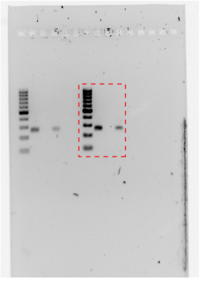
Extended Data Fig. 1o

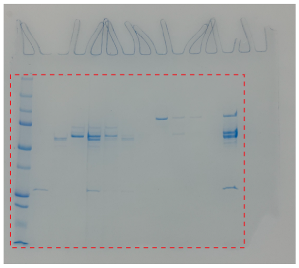
Extended Data Fig. 2c

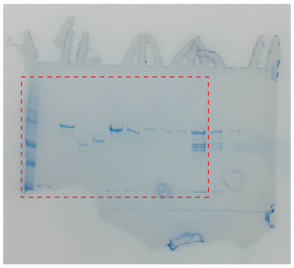
Extended Data Fig. 2f

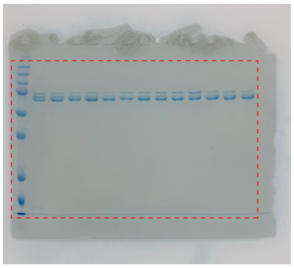
Extended Data Fig. 2c

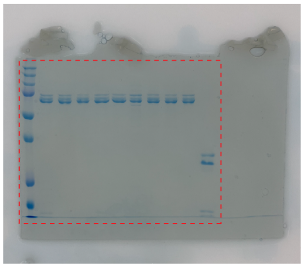
Extended Data Fig. 2c

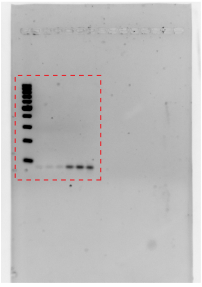
Extended Data Fig. 7a

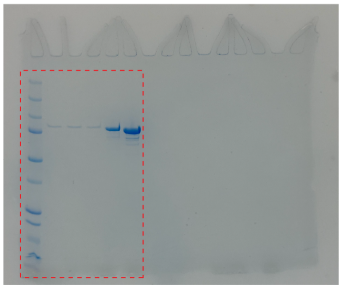
Extended Data Fig. 7a

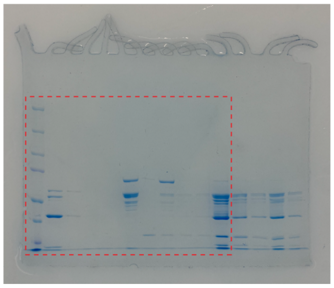
Extended Data Fig. 8a

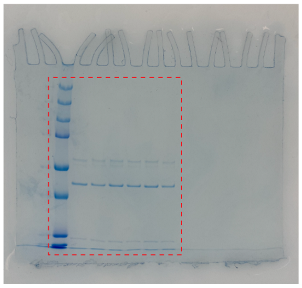
Extended Data Fig. 8k

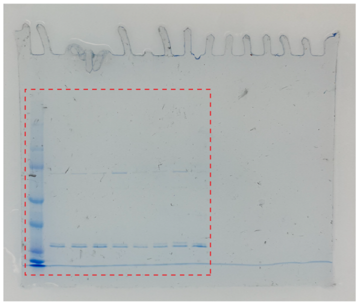
Extended Data Fig. 8l

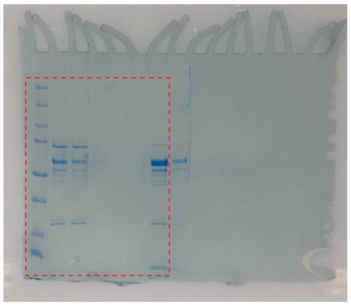
Extended Data Fig. 9a

**Supplementary Figure 1 | Source data for electrophoretic separation experiments.**

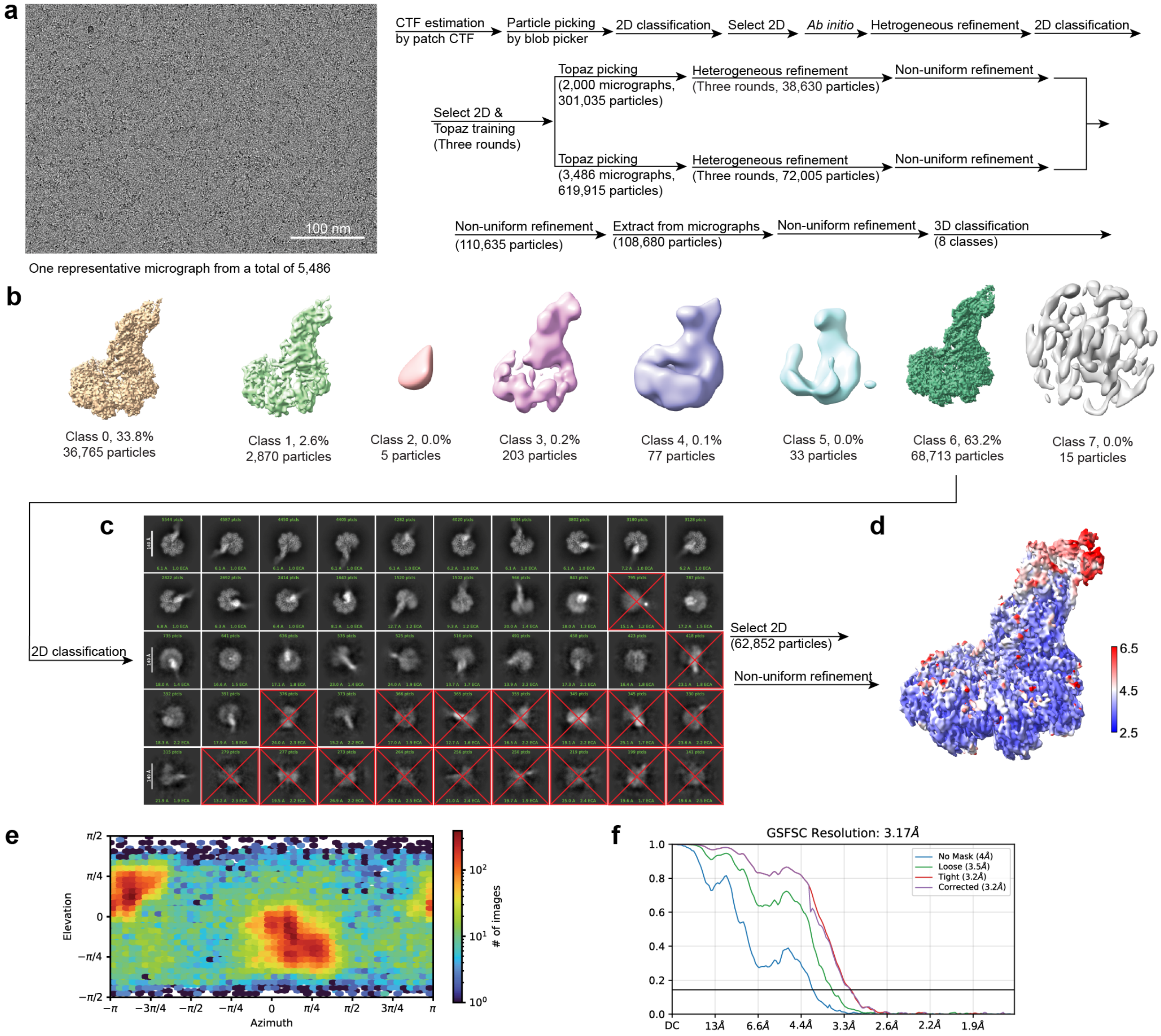


**Supplementary Figure 2 | Cryo-EM analysis of the $TnsC_7$-$TnsD_1$-$DNA_1$ complex showing preferred orientation.**

**a,** Representative raw cryo-EM micrograph from a dataset of 5,486 micrographs.

**b,** Three-dimensional classification of particles after non-uniform refinement.

**c,** Representative 2D class averages of selected particles from Class 6.

**d,** Local-resolution map of the $TnsC_7$-$TnsD_1$-$DNA_1$ reconstruction.

**e,** Viewing-direction distribution of particles used for the reconstruction, showing preferred orientation.

**f,** Gold-standard FSC curve for the $TnsC_7$-$TnsD_1$-$DNA_1$ reconstruction.

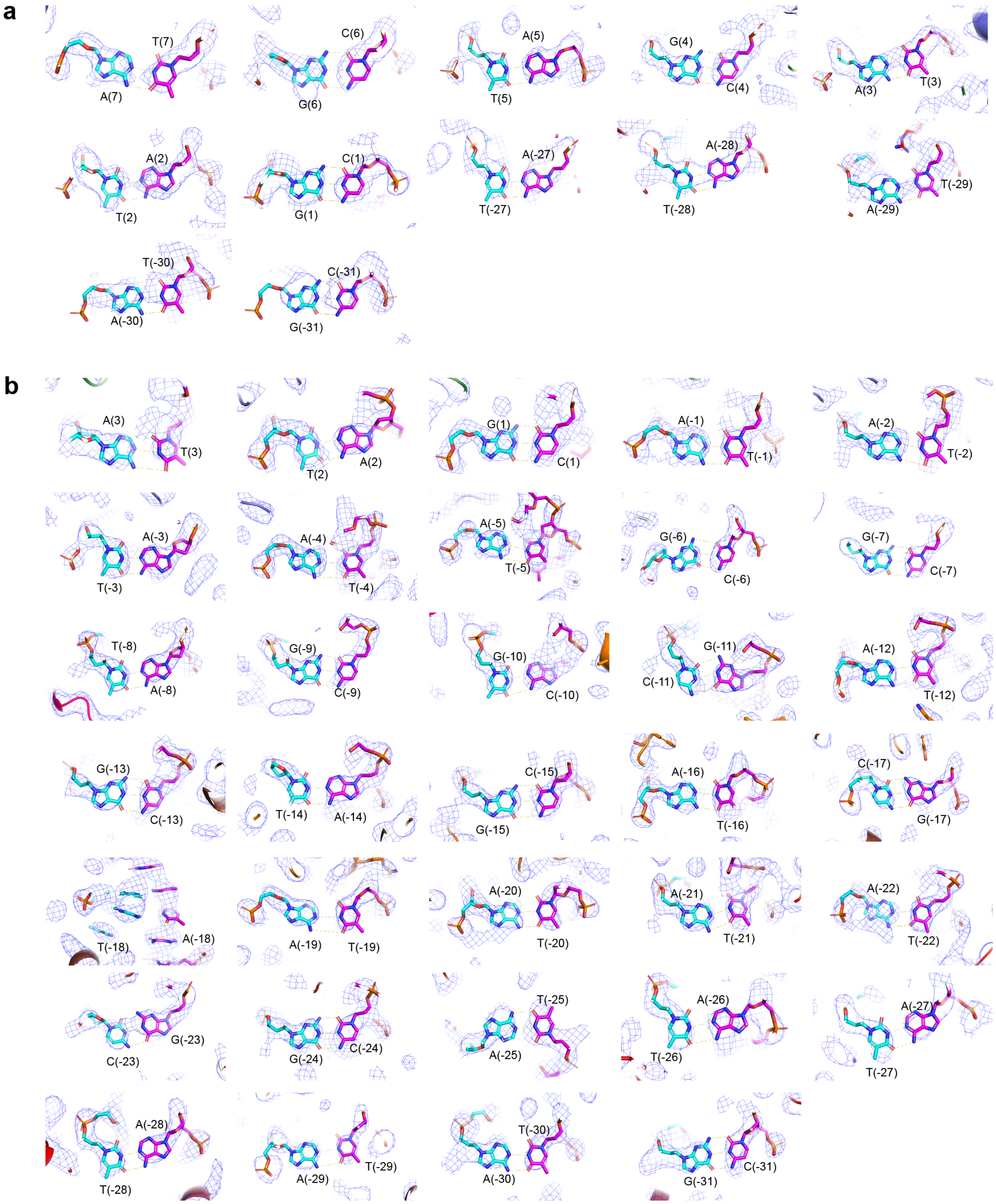


**Supplementary Figure 3 | DNA model fitting into cryo-EM density in TnsD–TnsC–DNA assemblies.**

**a,b,** Representative density and model views of individual base pairs in the $TnsC_3$-$TnsD_1$-$DNA_1$ (**a**) and $TnsC_7$-$TnsD_1$-$DNA_1$ (**b**) structures.

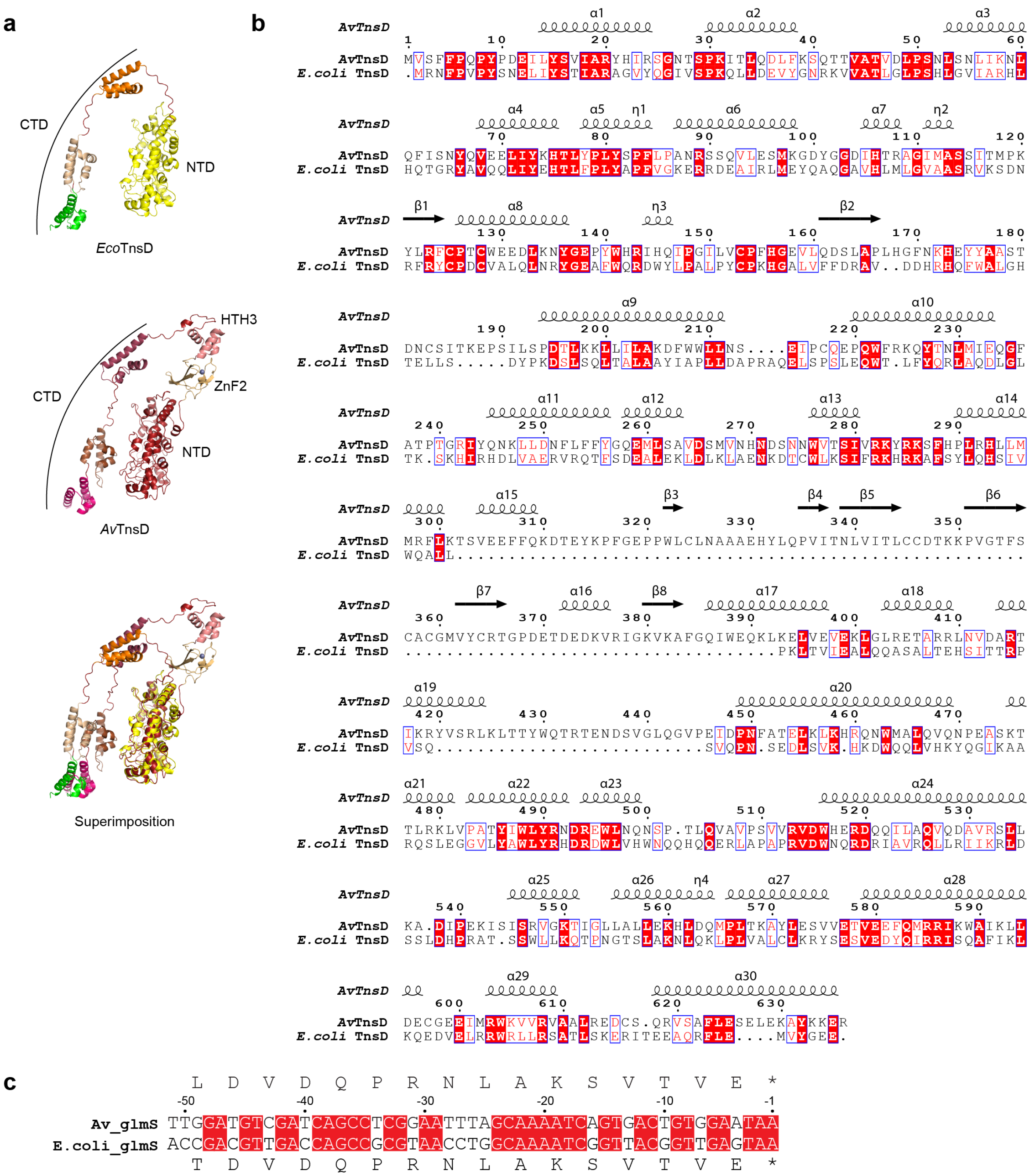


**Supplementary Figure 4 | Comparison of *Av*TnsD with Tn7 TnsD.**

**a,** AlphaFold-predicted structure of *E. coli* Tn7 TnsD (*Eco*TnsD), the structure of *Av*TnsD from the $TnsC_7$–$TnsD_1$–$DNA_1$ complex, and their structural superposition.

**b,** Sequence alignment of *Av*TnsD and *Eco*TnsD.

**c,** Sequence alignment of the 51-bp regions at the 3′ ends of *glmS* from *Anabaena variabilis* and *E. coli*.

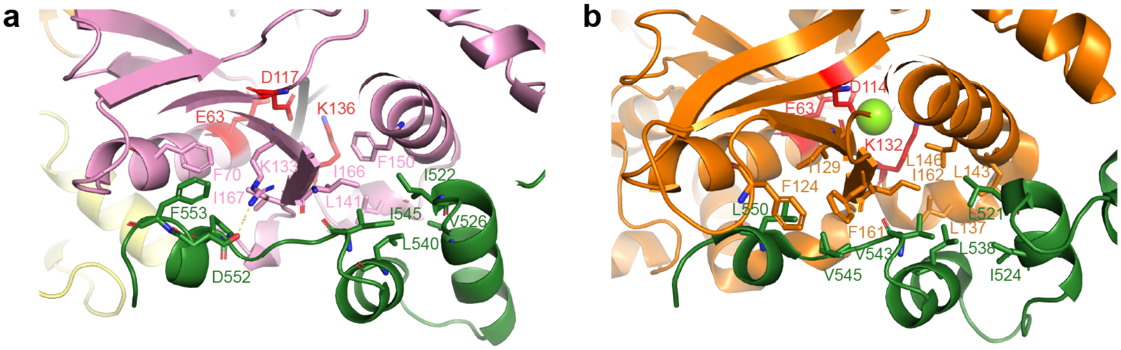


**Supplementary Figure 5 | Conserved TnsA–TnsC interactions in *Av*CAST and Tn7.**

**a,** Detailed interactions between *Av*TnsA and the C-terminal region of *Av*TnsC (residues 513–556) in the *Av*CAST STC.

**b,** Detailed interactions between *Eco*TnsA and the C-terminal region of *Eco*TnsC (residues 504–555) in the crystal structure of the *Eco*TnsA–TnsC complex (PDB: 1T0F).

**Supplementary Table 1 | DNA oligos used in this study.**

| DNA oligo | Sequence | Purpose |
|---|---|---|
| pRSFDuet_MCS2_Cas5_F | GGAGATATACATATGACGACAATTGCCTTAAAAGTAGAAGTTCCC | Amplification of Cas5 gene |
| pRSFDuet-1_MCS2_Cas5_R | AGCAGCCTAGGTTAACTAATTACTTTGAACTGAAGTCC | Amplification of Cas5 gene |
| pRSFDuet-1_MCS1_F | TGCTTAAGTCGAACAGAAAGTAATCG | Linearization of pRSFDuet-1 vector for Cas5 insertion |
| pRSFDuet-1_MCS1_R | GGTATATCTCCTTATTAAAGTTAAACAAAATTATTTCTACAGGGG | Linearization of pRSFDuet-1 vector for Cas5 insertion |
| pCDFDuet-1_MCS2_Cas6_F | GAAGGAGATATACATATGATTATTGATAGTAACTTAGTTACGCAGC | Amplification of Cas6 gene |
| pCDFDuet-1_MCS2_Cas6_R | AGCAGCCTAGGTTAACTATTGCCTTTCCTTAATAGG | Amplification of Cas6 gene |
| pCDFDuet-1_MCS2_F | TTAACCTAGGCTGCTGCCACC | Linearization of pCDFDuet-1 vector for Cas6 insertion |
| pCDFDuet-1_MCS2_R | ATGTATATCTCCTTCTTATACTTAACTAATATACTAAGATGGGG | Linearization of pCDFDuet-1 vector for Cas6 insertion |
| Cas7_pRSFDuet-1_F | ATAAGGAGATATACCATGTTTCATCTGTTTGGTAATATTTTGAC | Amplification of Cas7 gene |
| Cas7_pRSFDuet-1_R | TGTTCGACTTAAGCATCATGAACTCCTTAACGACAAC | Amplification of Cas7 gene |
| pRSFDuet-1_MCS1_F | TGCTTAAGTCGAACAGAAAGTAATCG | Linearization of pRSFDuet-1 vector for Cas7 insertion |
| pRSFDuet-1_MCS1_R | GGTATATCTCCTTATTAAAGTTAAACAAAATTATTTCTACAGGGG | Linearization of pRSFDuet-1 vector for Cas7 insertion |
| Cas8_2_F | CACCATCATCACCACATGAAGAGAGCAGATAACCCC | Amplification of Cas8 gene |

| | | |
|---|---|---|
| Cas8_2_R | TGTTCGACTTAAGCATTATTCAGTATTTAGC TCCGATATTTGTG | Amplification of Cas8 gene |
| pETDuet-2_F | TGCTTAAGTCGAACAGAAAGTAATC | Linearization of pETDuet-1 vector for Cas8 insertion |
| pETDuet-2_R | GTGGTGATGATGGTGATGGC | Linearization of pETDuet-1 vector for Cas8 insertion |
| CRISPR_pCDFDuet-1_MCS1_F | AGGAGATATACCATGGTGCTTTAACATTAG ATGTCGTTAGG | Amplification of CRISPR array with one spacer and two direct repeats |
| CRISPR_PCDFDuet-1_MCS1_R | GCTCGAATTCGGATCTGCTCAACGCCTAAC G | Amplification of CRISPR array with one spacer and two direct repeats |
| pCDFDuet-1_MCS1_F | GATCCGAATTCGAGCTCGGCGCG | Linearization of pCDFDuet-1 vector for CRISPR array insertion |
| pCDFDuet-1_MCS1_R | CATGGTATATCTCCTTATTAAAGTTAAACAA AATTATTTCTACAGGGG | Linearization of pCDFDuet-1 vector for CRISPR array insertion |
| AvTniQ_F | CAGATTGGTGGATCCCTTAGTTTTTCCCAA CTCTGTATCCC | Amplification of the AvTniQ gene |
| AvTniQ_R | CTCGAGTGCGGCCGCTTATGACTGAAAGAA CTCTTC | Amplification of the AvTniQ gene |
| TnsB_Backbone_F | GCGGCCGCACTCGAGGCC | Linearization of pTwinStrep-SUMO vector for AvTniQ insertion |
| TnsB_Backbone_R | GGATCCACCAATCTGTTCTCTGTGAGCC | Linearization of pTwinStrep-SUMO vector for AvTniQ insertion |
| 12k_backbone_f | TGTATGGTGCACTCTCAGTAC | Construction of pDonor_KanR plasmid |
| 12k_backbone_r | CTTAATTAGCTGAGCTTGGACTCC | Construction of pDonor_KanR plasmid |
| AvLE_RS_F | GCTCAGCTAATTAAGCCGGGATATTTACGTT TTGTAATCA | Construction of pDonor_KanR plasmid |

| | | |
|---|---|---|
| AvLE_RS_R | AGAGTGCACCATACATGTAGACAAAGAATTCGGTTAAACA | Construction of pDonor_KanR plasmid |
| 12k_backbone_2_F | AGGGGGATCCACTAGTGAGC | Construction of pDonor_KanR plasmid |
| 12k_backbone_2_R | TTACTTTCCGCTGCATAACCC | Construction of pDonor_KanR plasmid |
| AvRE_RS_F | TGCAGCGGAAAGTAATGTGGAGGAATAATTCGGTTCAAC | Construction of pDonor_KanR plasmid |
| AvRE_RS_R | CTAGTGGATCCCCCTCCATTCCCTTCTACCCCTAACC | Construction of pDonor_KanR plasmid |
| pTarget_AvPSP1_f | GATACGTAGATCAATGAGAAGTCATTTAATAAGGCCACTGT | Construction of pTarget plasmid |
| pTarget_AvPSP1_r | CCTTATTAAATGACTTCTCATTGATCTACGTATCGTAGATACT | Construction of pTarget plasmid |
| pTarget_lin_f | GTAGTGATCTTATTTCATTATGGTGAAAGTTGGAACC | Construction of pTarget plasmid |
| pTarget_lin_r | CGGGCGTATTTTTTGAGTTATCGAGATTTTCAGG | Construction of pTarget plasmid |
| glms_50_1 | CAAAAAATACGCCCGTGGATGTCGATCAGCCTCGGAATTTAGCAAAATCAGTGACTGTGGAATAAGTAGTGATCTTATTT | Construction of pTarget plasmid |
| glms_50_2 | AAATAAGATCACTACTTATTCCACAGTCACTGATTTTGCTAAATTCCGAGGCTGATCGACATCCACGGGCGTATTTTTTG | Construction of pTarget plasmid |
| glmS_Downstream_qPCR | GCACGTAAGAGGTTCCAACT | qPCR primer for TnsD-guided transposition |
| Av_LE | ACCTTTATCAGTAATGGATTCAAAC | qPCR primer for TnsD-guided transposition |
| Av_LE_probe | ACAGTTCGAATATTTAATCTGCCGTTGTCG | qPCR probe |
| PSP1_r1 | GGAAATCGCTGCGTATGTG | qPCR primer for RNA-guided transposition |
| Av_LE_q3 | CACAGCAGAACTTTCTCAAATAATTTTG | qPCR primer for RNA-guided transposition |
| TC_f | CGACAGCATCGCCAGTCACTATG | pTarget control forward primer |
| TC_r | CAAGTAGCGAAGCGAGCAGGAC | pTarget control reverse Primer |

| | | |
|---|---|---|
| TC_probe | TGCGTTGATGCAATTTCTATGCGCACCCGT | qPCR probe for pTarget control |
| RE30_F | acaatataagggggatccactagtgag | RE Truncation Primer |
| RE30_R | tccccttatattgttgaaccgaattattcc | RE Truncation Primer |
| RE40_F | attcggttagggggatccactagtgag | RE Truncation Primer |
| RE40_R | tcccctaaccgaattatatattgttgaacc | RE Truncation Primer |
| RE50_F | acaattaaagggggatccactagtgagc | RE Truncation Primer |
| RE50_R | tccccctttaattgttgaaccgaattatatattg | RE Truncation Primer |
| RE70_F | acaataacagggggatccactagtgagc | RE Truncation Primer |
| RE70_R | tccccctgttattgttcaatcgttttattaattg | RE Truncation Primer |
| RE80_F | ctaaaggaagggggatccactagtgagc | RE Truncation Primer |
| RE80_R | tccccttcctttagaggttattgttcaatc | RE Truncation Primer |
| RE100_F | caagtaaaagggggatccactagtgagc | RE Truncation Primer |
| RE100_R | tcccccttttacttggtacaatcgtactcc | RE Truncation Primer |
| LE30_F | taattaagataattgtttaaccgaattctttgtc | LE Truncation Primer |
| LE30_R | caattatcttaattagctgagcttggactc | LE Truncation Primer |
| LE90_F | taattaaggctgaaattgttcaatcgttttattttatcg | LE Truncation Primer |
| LE90_R | tttcagccttaattagctgagcttggactc | LE Truncation Primer |
| LE160_F | taattaagggattcaaacaattgtttaaccg | LE Truncation Primer |
| LE160_R | tgaatcccttaattagctgagcttggac | LE Truncation Primer |
| L1F_dele_F | gaactgtatgtctacatgtatggtgcactc | LE Truncation Primer |

| | | |
|---|---|---|
| L1F_dele_R | gtagacatacagttcgaatatttaatctgc | LE Truncation Primer |
| L2F_dele_F | gttaagctttatcgacaacggcagattaaatattcg | LE Truncation Primer |
| L2F_dele_R | tcgataaagcttaacctctatggtaaacttg | LE Truncation Primer |
| L3F_dele_F | ggattcaatgagatttttactaaaaatcaagtttacc | LE Truncation Primer |
| L3F_dele_R | aatctcattgaatccattactgataaagg | LE Truncation Primer |
| L12F_dele_F | gttaagcttgtctacatgtatggtgcactc | LE Truncation Primer |
| L12F_dele_R | gtagacaagcttaacctctatggtaaacttg | LE Truncation Primer |
| L23F_dele_F | ggattcaattatcgacaacggcagattaaatattcg | LE Truncation Primer |
| L23F_dele_R | tcgataattgaatccattactgataaaggttttgag | LE Truncation Primer |
| L123F_dele_F | ggattcaatgtctacatgtatggtgcactc | LE Truncation Primer |
| L123F_dele_R | gtagacattgaatccattactgataaaggttttgag | LE Truncation Primer |
| REAssay_F | tcagtgactgtggaataagtagtg | RE Truncation assay forward primer |
| REAssay_R | tcctattccgaagttcctattctc | RE Truncation assay reverse primer |
| LRAssay_F | atccagatggagttctgaggtc | LE Truncation assay forward primer |
| LETAssay_R | gacgttgatcggcacgtaag | LE Truncation assay reverse primer |
| R285A_r | ATGGAAAGATTTCGCATATTTCCTGACGAT GCT | TnsD R285A mutation |
| K349A_f | TGCTGCGATACAGCGAAGCCTGTCGGTACT T | TnsD K349A mutation |
| K349A_r | ACCGACAGGCTTCGCTGTATCGCAGCATAA AGT | TnsD K349A mutation |
| K380A_f | GTTCGCATTGGTGCGGTCAAGGCATTCGGT C | TnsD K380A mutation |

| | | |
|---|---|---|
| K380A_r | GAATGCCTTGACCGCACCAATGCGAACTTTGTC | TnsD K380A mutation |
| R404A_f | AAGTTGGGTTTGGCGGAGACAGCCAGAAGATT | TnsD R404A mutation |
| R404A_r | TCTGGCTGTCTCCGCCAAACCCAACTTTTCAAC | TnsD R404A mutation |
| R415A_f | AATGTTGATGCCGCGACTATCAAACGTTATGTAT | TnsD R415A mutation |
| R415A_r | ACGTTTGATAGTCGCGGCATCAACATTTAATCT | TnsD R415A mutation |
| R479A_f | AAGACAACTCTGGCGAAACTAGTACCAGCCACC | TnsD R479A mutation |
| R479A_r | TGGTACTAGTTTCGCCAGAGTTGTCTTTGAAGC | TnsD R479A mutation |
| R491A_f | ATTTGGCTATACGCGAATGACAGAGAATGGCTC | TnsD R491A mutation |
| R491A_r | TTCTCTGTCATTCGCGTATAGCCAAATATAGGT | TnsD R491A mutation |
| R513A_f | CCTTCCGTTGTGGCGGTTGATTGGCATG | TnsD R513A mutation |
| R513A_r | ATGCCAATCAACCGCCACAACGGAAGGAAC | TnsD R513A mutation |
| K550A_f | AGTAGGGTAGGGGCGACTATCGGCTTACTTGC | TnsD K550A mutation |
| K550A_r | TAAGCCGATAGTCGCCCCTACCCTACTAATAG | TnsD K550A mutation |
| K560A_f | GCATTACTCGAAGCGCATCTCGACCAGATGCCT | TnsD K560A mutation |
| K560A_r | CTGGTCGAGATGCGCTTCGAGTAATGCAAGTAAG | TnsD K560A mutation |
| R608A_f | TGGAAGGTAGTCGCGGTTGCTGCTTTGCGAGAG | TnsD R608A mutation |
| R608A_r | CAAAGCAGCAACCGCGACTACCTTCCAGCGCAT | TnsD R608A mutation |
| AvTnsD_dele_f | GAATTTTTTCAAAAATGAGCGGCCGCACTCGAGG | TnsD NTD mutation |
| AvTnsD_dele_r | TTTTTGAAAAAATTCCTCTACTGATGTTTTCAAAAAACGC | TnsD NTD mutation |
| Cas8_H117A_f | GCCACTTTTACACAAGCGAATAAATTTTTAAGTCT | Cas8 H117A mutation |

| | | |
|---|---|---|
| Cas8_H117A_r | AAAAAATTTATTCGCTTGTGTAAAAGTGGCTGT | Cas8 H117A mutation |
| Cas8_Q295A_f | GCTTGGTCAGAAGCGCAAAAAACTCGTGCAG | Cas8 Q295A mutation |
| Cas8_Q295A_r | ACGAGTTTTTTGCGCTTCTGACCAAGCTTTTGTAC | Cas8 Q295A mutation |
| Cas8_Q296A_f | TGGTCAGAACAGGCGAAAACTCGTGCAGAAATAG | Cas8 Q296A mutation |
| Cas8_Q296A_r | TGCACGAGTTTTCGCCTGTTCTGACCAAGCTTT | Cas8 Q296A mutation |
| Cas5_R75A_f | CGCACCTTTCACGCGTTTAAGACGAAGAATATCCAT | Cas5 R75A mutation |
| Cas5_R75A_r | ATTCTTCGTCTTAAACGCGTGAAAGGTGCGAATAACAGT | Cas5 R75A mutation |
| Cas5_K77A_f | CTTTCACCGATTTGCGACGAAGAATATCCATGACT | Cas5 K77A mutation |
| Cas5_K77A_r | GGATATTCTTCGTCGCAAATCGGTGAAAGGTGCG | Cas5 K77A mutation |
| TniQ_Y280A_f | AATCAAGGTAAAGCGTTTTCTCACTGCTTGTTT | TniQ Y280A mutation |
| TniQ_Y280A_r | GCAGTGAGAAAACGCTTTACCTTGATTTTGGGT | TniQ Y280A mutation |
| TniQ_H283A_f | GTAAATATTTTTCTGCGTGCTTGTTTGGATGCGAC | TniQ H283A mutation |
| TniQ_H283A_r | GCATCCAAACAAGCACGCAGAAAAATATTTACCTTG | TniQ H283A mutation |
| TniQ_D289A_f | TTGTTTGGATGCGCGTTAAACCCAGCAATTGAC | TniQ D289A mutation |
| TniQ_D289A_r | TGCTGGGTTTAACGCGCATCCAAACAAGCAGTG | TniQ D289A mutation |
| TniQ_L290R_f | TTTGGATGCGACCGTAACCCAGCAATTGACCGC | TniQ L290R mutation |
| TniQ_L290R_r | AATTGCTGGGTTACGGTCGCATCCAAACAAGCAG | TniQ L290R mutation |
| TniQ_F241A_f | CTGCTAAAAACCGCGGCTGGTGGCAAGTTTAAG | TniQ F241A mutation |
| TniQ_F241A_r | CTTGCCACCAGCCGCGGTTTTTAGCAGTCCTT | TniQ F241A mutation |

| | | |
|---|---|---|
| TniQ_F246A_f | GCTGGTGGCAAGGCGAAGTTTAATGATTGT GGT | TniQ F246A mutation |
| TniQ_F246A_r | ATCATTAAACTTCGCCTTGCCACCAGCCGCG GT | TniQ F246A mutation |
| 239-247_dele_f | TTTAATGATTGTGGTTTTGCCCAGCT | TniQ ΔK239–K248 mutation |
| 239-247_dele_r | TAGCAGTCCTTTATCCAATAAGTAACGC | TniQ ΔK239–K248 mutation |
| 239-247_GSA9aa_f | GATAAAGGACTGCTAGGTAGTGCCGGCTCT GCAGGTTCTGCCTTTAATGATTGTGGT | TniQ ΔK239–K248 mutation |
| 239-247_GSA9aa_r | ACCACAATCATTAAAGGCAGAACCTGCAGA GCCGGCACTACCTAGCAGTCCTTTATC | TniQ ΔK239–K248 mutation |
| 287-293_dele_f | ATTGACCGCATAACTCATATCTTAATG | TniQ ΔG287–A293 mutation |
| 287-293_dele_r | AAACAAGCAGTGAGAAAAATATTTACCT | TniQ ΔG287–A293 mutation |
| 287-293_GSA7aa_f | TCTCACTGCTTGTTTGGTAGTGCCGGCTCTG CAGGTATTGACCGCATAACT | TniQ ΔG287–A293 mutation |
| 287-293_GSA7aa_r | AGTTATGCGGTCAATACCTGCAGAGCCGGC ACTACCAAACAAGCAGTGAGA | TniQ ΔG287–A293 mutation |
| glmS_target_G(-6)T_f | TCAGTGACTGTGTAATAAGTAGTGATCTTAT TTCATTATGG | pTarget G(-6)T mutation |
| glmS_target_G(-6)T_r | TCACTACTTATTACACAGTCACTGATTTTGC TAAATTCC | pTarget G(-6)T mutation |
| glmS_target_C(-11)A_f | GCAAAATCAGTGAATGTGGAATAAGTAGTG ATC | pTarget C(-11)A mutation |
| glmS_target_C(-11)A_r | ACTTATTCCACATTCACTGATTTTGCTAAAT TCC | pTarget C(-11)A mutation |
| glmS_target_C(-17)A_f | ATTTAGCAAAATAAGTGACTGTGGAATAAG TAGTG | pTarget C(-17)A mutation |
| glmS_target_C(-17)A_r | TCCACAGTCACTTATTTTGCTAAATTCCGAG | pTarget C(-17)A mutation |
| glmS_target_T(-18)G_f | AATTTAGCAAAAGCAGTGACTGTGGAATAA GTAGTG | pTarget T(-18)G mutation |
| glmS_target_T(-18)G_r | CCACAGTCACTGCTTTTGCTAAATTCCGAGG | pTarget T(-18)G mutation |

| | | |
|---|---|---|
| glmS_target_G(-24)T_f | CCTCGGAATTTAT CAAAATCAGTGACTGTGG | pTarget G(-24)T mutation |
| glmS_target_G(-24)T_r | GTCACTGATTTTGA TAAATTCCGAGGCTGATC | pTarget G(-24)T mutation |
| glmS_target_A(-29)C_f | ATCAGCCTCGGACTTTAGCAAAATCAGTGA C | pTarget A(-29)C mutation |
| glmS_target_A(-29)C_r | GATTTTGCTAAAGTCCGAGGCTGATCGACA TC | pTarget A(-29)C mutation |
| glmS_target_G(-32)T_f | TCGATCAGCCTCTGAATTTAGCAAAATCAG TG | pTarget G(-32)T mutation |

### Supplementary Video

Supplementary Video 1 | Morphing of TnsD from the $TnsC_3$-$TnsD_1$-$DNA_1$ complex to the $TnsC_7$-$TnsD_1$-$DNA_1$ complex, with the TnsC heptamer from the latter shown in transparency. HTH4–6 domains are not resolved in the $TnsC_3$-$TnsD_1$-$DNA_1$ structure and are modeled for clarity.

Supplementary Video 2 | Morphing of *Av*Cascade-DNA-1, *Av*Cascade-DNA-2, and *Av*Cascade-TniQ-DNA structures.
