## Extended Data Tabel 1 for "DNA remodeling couples target recognition to directional transposition in a Tn7-like CAST"

### Extended Data Table 1 | Cryo-EM data collection, refinement and validation statistics

| | $TnsC_3$–$TnsD_1$–$DNA_1$ | $TnsC_4$–$TnsD_1$–$DNA_1$ | $TnsC_7$–$TnsD_1$–$DNA_1$ | STC | TnsB–DNA | *Av*Cascade–DNA-1 | *Av*Cascade–DNA-2 | *Av*Cascade–TniQ–DNA | TniQ-TnsC-DNA |
|---|---|---|---|---|---|---|---|---|---|
| **Data collection and processing** | | | | | | | | | |
| Magnification | 105,000 | 105,000 | 105,000 | 105,000 | 105,000 | 105,000 | 105,000 | 105,000 | 105,000 |
| Voltage (kV) | 300 | 300 | 300 | 300 | 300 | 300 | 300 | 300 | 300 |
| Electron exposure (e–/Å$^2$) | 59.5 | 59.5 | 59.5 | 59.5 | 59.5 | 59.5 | 59.5 | 59.5 | 59.5 |
| Defocus range (μm) | 0.8–2.0 | 0.8–2.0 | 0.8–2.0 | 0.8–2.0 | 0.8–2.0 | 0.8–2.0 | 0.8–2.0 | 0.8–2.0 | 0.8–2.0 |
| Pixel size (Å) | 0.822 | 0.822 | 0.822 | 0.822 | 0.822 | 0.822 | 0.822 | 0.822 | 0.822 |
| Symmetry imposed | C1 | C1 | C1 | C1 | C1 | C1 | C1 | C1 | C1 |
| Initial particle images (no.) | 788,816 | 788,816 | 787,659 | 1,851,842 | 295,815 | 1,404,457 | 1,404,457 | 1,404,457 | 332,402 |
| Final particle images (no.) | 91,633 | 11,240 | 166,543 | 51,307 | 53,796 | 194,480 | 49,799 | 42,834 | 22,162 |
| Map resolution (Å) | 3.08 | 3.82 | 3.23 | 3.19 | 3.9 | 2.75 | 3.01 | 3.16 | 3.56 |
| FSC threshold | 0.143 | 0.143 | 0.143 | 0.143 | 0.143 | 0.143 | 0.143 | 0.143 | 0.143 |
| Map resolution range (Å) | 2.7–50.1 | 3.2–61.2 | 1.75–47.0 | 2.8–52.3 | 3.1–64.4 | 2.3–36.1 | 2.6–47.8 | 1.7–52.3 | 1.7–61.3 |
| **Refinement** | | | | | | | | | |
| Initial model used | AF Models | AF Models | AF Models | AF Models | AF Models | AF Models | AF Models | AF Models | AF Models |
| Model resolution (Å) | 3.2 | 3.9 | 3.8 | 3.5 | 8.6 | 2.9 | 3.3 | 3.0 | 3.9 |
| FSC threshold | 0.5 | 0.5 | 0.5 | 0.5 | 0.5 | 0.5 | 0.5 | 0.5 | 0.5 |
| Model resolution range (Å) | 2.4–50 | 2.4–50 | 2.4–50 | 2.4–50 | 2.4–50 | 2.4–50 | 2.4–50 | 2.4–50 | 2.4–50 |
| Map sharpening *B* factor (Å$^2$) | –75.2 | –18.7 | –85.7 | –51.8 | –74.9 | –81.8 | –54.4 | –47.7 | –21.8 |
| Model composition | | | | | | | | | |
| Non-hydrogen atoms | 15546 | 18746 | 30123 | 35211 | 16611 | 28865 | 29313 | 29767 | 26888 |
| Protein residues | 1617 | 2009 | 3387 | 3587 | 1724 | 3261 | 3257 | 3312 | 3157 |
| Nucleotides | 114 | 114 | 120 | 280 | 116 | 129 | 153 | 150 | 52 |
| Ligands | 3 | 4 | 7 | 0 | 0 | 0 | 0 | 0 | 7 |
| Mg | 3 | 4 | 7 | 4 | 0 | 0 | 0 | 0 | 7 |
| Zn | 2 | 2 | 2 | 0 | 0 | 0 | 0 | 1 | 1 |
| *B* factors (Å$^2$) | | | | | | | | | |
| Protein | 144.00 | 140.00 | 206.30 | 220.26 | 421.62 | 133.24 | 160.05 | 75.85 | 134.77 |
| Nucleic acids | 285.67 | 255.70 | 229.23 | 256.57 | 227.73 | 187.49 | 203.41 | 101.00 | 187.90 |
| Ligand | 106.35 | 108.67 | 198.37 | 109.59 | - | - | - | 136.59 | 123.15 |
| R.m.s. deviations | | | | | | | | | |
| Bond lengths (Å) | 0.005 | 0.003 | 0.005 | 0.003 | 0.004 | 0.005 | 0.005 | 0.005 | 0.003 |
| Bond angles (°) | 0.898 | 0.625 | 0.962 | 0.536 | 0.835 | 0.953 | 0.936 | 0.931 | 0.702 |
| Validation | | | | | | | | | |
| MolProbity score | 2.19 | 2.22 | 2.35 | 1.96 | 2.71 | 1.91 | 1.90 | 2.23 | 2.46 |
| Clashscore | 8.39 | 10.56 | 10.64 | 8.75 | 23.53 | 9.22 | 9.65 | 8.98 | 12.66 |
| Poor rotamers (%) | 2.36 | 2.13 | 2.99 | 2.35 | 2.94 | 1.59 | 1.52 | 3.85 | 3.07 |
| Ramachandran plot | | | | | | | | | |
| Favored (%) | 92.98 | 93.45 | 93.24 | 96.66 | 91.94 | 96.09 | 96.24 | 95.62 | 92.39 |
| Allowed (%) | 6.40 | 6.30 | 6.20 | 3.15 | 7.65 | 3.79 | 3.61 | 4.10 | 7.26 |
| Disallowed (%) | 0.62 | 0.25 | 0.56 | 0.20 | 0.41 | 0.12 | 0.16 | 0.28 | 0.35 |
